# Rational design of potent small molecule SMARCA2/A4 (BRM/BRG1) degraders acting via the recruitment of FBXO22

**DOI:** 10.1101/2025.03.05.641484

**Authors:** Elisia Villemure, Tom Januario, Mingshuo Zeng, Hanna G. Budayeva, Benjamin T. Walters, Aaron Lictao, Ke Sherry Li, Xiaofen Ye, Caroline L. Gilchrist, Bridget Hoag, Nicholas F. Endres, Peter L. Hsu, John Chan, Tommy K. Cheung, Michael R. Costa, Jean-Philippe Fortin, Noriko Ishisoko, Brett M. Babin, Joyce Liu, Joachim Rudolph, Robert L. Yauch

## Abstract

Target-anchored monovalent degraders are more drug-like than their bivalent counterparts, Proteolysis Targeting Chimeras (PROTACs), while offering greater target specificity control than the E3 ligase-anchored monovalent degraders, also known as molecular glues. However, their discovery has typically been serendipitous, and the rules governing their identification remain unclear. This study focused on the intentional discovery of SMARCA2/A4 monovalent degraders using a library based on SMARCA2/A4 bromodomain-binding ligands. Compound **G-6599** emerged as a lead candidate, showing exceptional degradation potency and specificity for SMARCA2/A4. Mechanistic studies revealed that **G-6599** operates through the ubiquitin-proteasome pathway and the E3 ligase FBXO22. **G-6599** was shown to promote ternary complex formation between SMARCA2 and FBXO22 involving covalent conjugation to a cysteine residue on the latter. Unlike other recently identified FBXO22-dependent degraders, it does not require biotransformation. The selective degradation ability of **G-6599**, along with its unique mechanism, highlights the therapeutic potential of target-anchored monovalent degraders.

## Introduction

Targeted protein degradation is a transformative paradigm in drug discovery that offers major promise as a therapeutic approach^1,2^. It relies on removing the disease-associated protein instead of solely inhibiting its function. Multiple protein degradation modalities are currently being investigated in the clinic and in pre-clinical research settings^3^. Bivalent degraders, also known as PROTACs (Proteolysis Targeting Chimeras), are prominently represented in the pipeline of degrader molecules undergoing clinical evaluation. They are heterobifunctional compounds that recruit a protein of interest to an E3 ligase, leading to the ubiquitination and proteasomal degradation of the former. The great majority of the bivalent degraders currently investigated in the clinic operate through the E3 ligase Cereblon (CRBN) and are in many cases orally bioavailable. While we are gaining a better understanding of how to achieve oral bioavailability with CRBN-based bivalent degraders^4,5^, challenges still persist.

E3 ligase-anchored monovalent degraders, such as the immunomodulatory drugs (IMiDs), which induce proximity of various targets to CRBN, are well-proven drugs that overcome some of these challenges^6–8^. They can enable the recruitment and degradation of poorly tractable proteins that do not necessarily possess a suitable binding pocket for small molecule ligands. However, the spectrum of proteins that can be recruited to CRBN or other E3 ligases by molecular glues and the rules governing the formation of a productive ternary complex still have yet to be fully understood.

Monovalent degraders binding to the protein of interest provide an interesting alternative with greater control over target specificity and more favorable drug-like properties than their bivalent counterparts. However, except for very recent examples^9–16^, most target-anchored monovalent degraders have been found serendipitously^17–21^. It would be highly desirable to use the ligand of a protein of interest as a starting point to rationally design monovalent degraders. An approach toward this would be to elaborate the solvent-exposed region, thereby creating a novel neo-surface that could promote the recruitment of an E3 ligase. Monovalent degraders would be especially beneficial for proteins with existing ligands without a known function. The SWitch/Sucrose nonfermentable (SWI/SNF) complex ATPases, SMARCA2 and SMARCA4, fall in this category, as ligands that were discovered to bind to their bromodomains were found to be devoid of functional activity in relevant cancer cell lines^22,23^.

The proteins SMARCA2 and SMARCA4 represent attractive oncology targets. In particular, tumors derived from several lineages exhibit exquisite sensitivities to pan-inhibition of SMARCA2 and SMARCA4, including androgen receptor (AR) -dependent prostate cancers, uveal melanoma and acute myeloid leukemias^24^. In addition, inhibition of SMARCA2 in cancers that harbor loss-of-function mutations in SMARCA4 may represent a novel synthetic lethal strategy to treat such cancer types^22,25,26^. Both ATPase inhibitors and PROTACs targeting the bromodomain of SMARCA2 and SMARCA4 have been reported, some of which are under clinical evaluation,^27–32^ underscoring the strong therapeutic potential of targeting these proteins.

In this work, we identified novel monovalent degraders of SMARCA2/A4 that function through the recruitment of the F-box only protein 22 cullin-RING ligase (FBXO22 CRL) complex. These monovalent degraders were intentionally designed from SMARCA2/A4 bromodomain ligands and imparted potent degradation activity and anti-proliferative effects in AR-dependent prostate cancer cell lines.

## Results

### Identification of SMARCA2/A4 monovalent degraders

Our research campaign for SMARCA2/A4 monovalent degraders began with the screening of a library of compounds for their ability to deplete SMARCA2 levels in SW1573 cells as measured by an immunofluorescence assay. Our collection of small molecules was designed based on reported SMARCA2/A4 bromodomain ligands^33,34^, to which a diverse set of chemical moieties was appended at the solvent-exposed exit vector. We reasoned that these moieties could create a new neo-surface that may lead to recruitment of an E3 ligase and promote ubiquitination and proteasomal degradation of SMARCA2/A4. These entities were selected based on chemical diversity to cover a broad spectrum of charge, polarity and flexibility and were inspired by features of other reported small molecule degraders^17–21^. Upon screening of the library, using a SMARCA2 PROTAC as positive control, we identified two compounds, **2** and **3**, that imparted significant reduction of SMARCA2 nuclear intensity levels at a single concentration (Fig. 1a, Extended Data Fig. 1c). Additional studies confirmed dose dependency and rescue of degradation by a SMARCA2/A4 bromodomain ligand competition using parent ligand **1**, without associated cytotoxicity at the tested concentrations (Extended Data Fig. 1a, b, c). Compared to the parent ligand, monovalent degraders **2** and **3** displayed improved target engagement in a SMARCA2 NanoBRET^®^ assay in permeabilized cells and demonstrated moderate protein degradation with DC_50_ values of 13 nM and 18 nM, respectively, and a D_max_ of 38% in both cases (Table 1, Supplementary Table 1).

**Fig. 1.**
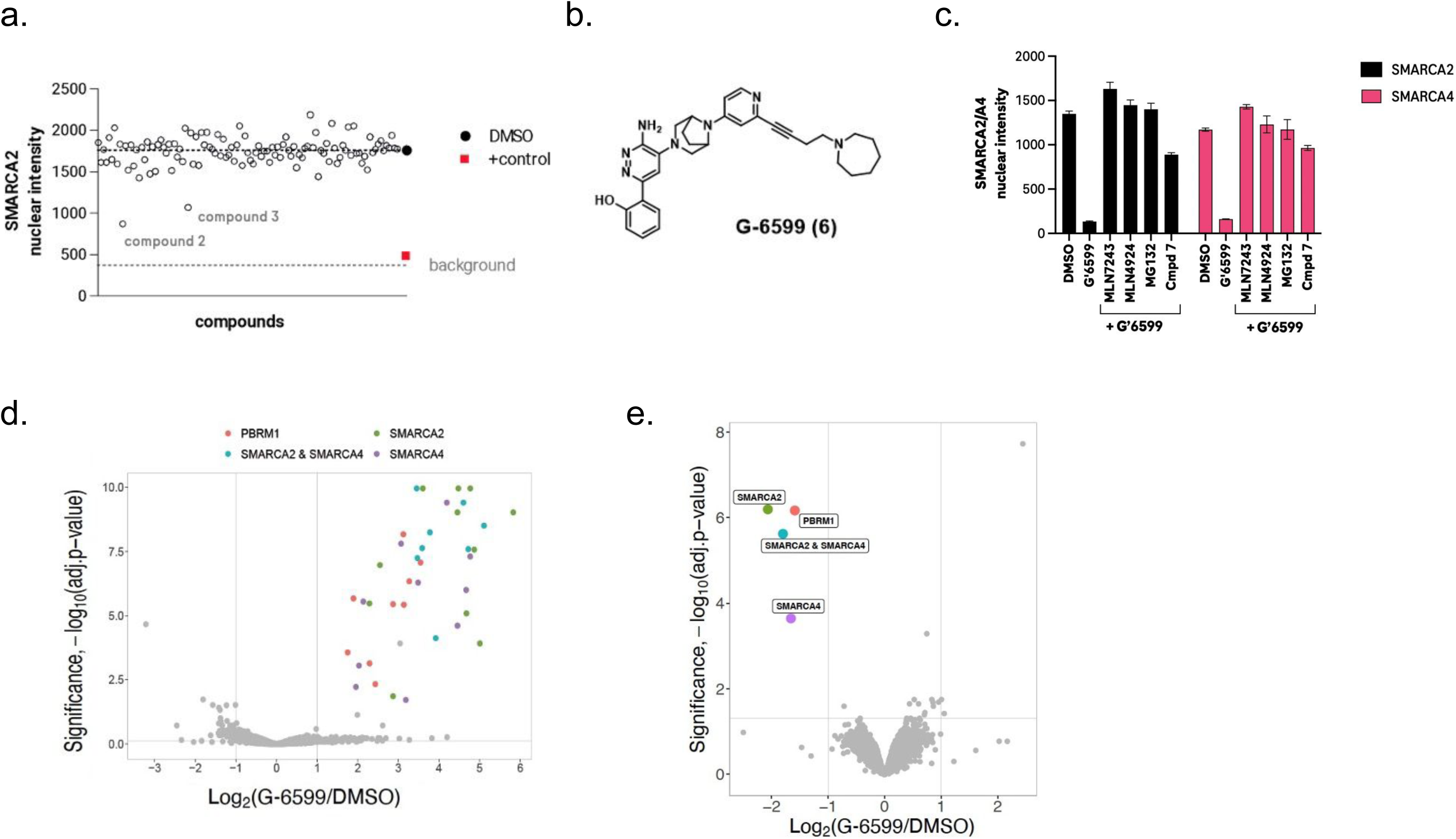
Identification of monovalent degraders of SMARCA2/A4. **a,** Immunofluorescence-based screen for degraders of SMARCA2 in SW1573 cells. A SMARCA2 PROTAC served as a positive control. The dotted line represents the background signal control without primary antibody. **b**, Chemical structure of potent monovalent degrader, **G-6599**. **c,** Pretreatment of SW1573 cells with 100-fold molar excess of the respective inhibitors (10 µM) can block **G-6599**-mediated degradation of SMARCA2 and SMARCA4, as detected by immunofluorescence. **d,** Global ubiquitylome changes assessed by di-glycine remnant mass spectrometry profiling in SW1573 cells following 30 min treatment with 100 nM **G-6599**. Peptide mapping to SMARCA2 (UniProtKB P-51531-2), SMARCA4 (UniProtKB P-51532-1) or PBRM1 (UniprotKB Q86U86) are highlighted in color. Ubiquitinated sites shared between SMARCA2 and SMARCA4 are shown in blue. n=9195 unique ubiquitinated sites were identified and presented as log2 fold change relative to DMSO control cultures. **e,** Global proteome assessed by mass spectrometry following 3 h treatment with **G-6599** (100 nM) in SW1573 cells. Data is presented as log2 fold change relative to DMSO control cultures. n=9717 proteins were quantified.

**Table 1.** Optimization of SMARCA2/A4 monovalent degraders.

| Compound | SMARCA2<br>DC<sub>50</sub> (nM) | SMARCA2<br>D<sub>max</sub> (%) | SMARCA4<br>DC<sub>50</sub> (nM) | SMARCA4<br>D<sub>max</sub> (%) |
| --- | --- | --- | --- | --- |
| **1** | >10,000 | -- | >10,000 | -- |
| **2** | 13 ± 40 | 38 ± 7.1 | 8.2 ± 2.4 | 50 ± 2.4 |
| **3** | 18 ± 25 | 38 ± 12 | 8.3 ± 5.3 | 43 ± 7.1 |
| **4** | 16 ± 6.5 | 51 ± 18 | 15 ± 9.9 | 25 ± 5.9 |
| **5** | 5.9 ± 2.1 | 82 ± 10 | 1.2 ± 0.75 | 99 ± 2.0 |
| **GNE-6599 (6)** | 0.018 ± 0.0092 | 97 ± 2.0 | 0.056 ± 0.062 | 95 ± 3.0 |

Following these promising results, we sought to improve degradation potency of our initial screening hits using a library expansion approach. We found that introduction of an alkyne spacer, as in compound **4**, can lead to improved SMARCA2 maximal degradation, while promoting weaker SMARCA4 degradation (Extended Data Fig. 1c, Table 1). We also observed that introducing rigidity in the linker using a cyclobutyl moiety to an analog of **2**, as in compound **5**, can lead to significant improvement in both SMARCA2 and SMARCA4 degradation, now achieving > 80% D_max_. After further optimization focusing on combining the alkyne spacer and a shorter amine linker, we identified **G-6599** (**6**), promoting exquisite degradation potency of both SMARCA2 and SMARCA4 with DC_50_ values of 18 pM and 56 pM, respectively (Fig. 1b, Table 1). **G-6599** also exhibited ≥ 95% maximal degradation of SMARCA2 and SMARCA4. The kinetics of degradation was comparable to A947, a previously published PROTAC, targeting SMARCA2 and SMARCA4, exhibiting > 90% degradation within 2 hours (Extended Data Fig. 1d). **G-6599** shared the alkyne spacer of compound **4**, while presenting an azepane tail at the solvent-exposed exit vector of the bromodomain binding pocket.

### Monovalent degrader G-6599 is specific to SMARCA2/A4 and operates through the ubiquitin-proteasome pathway

Mechanistic studies by immunofluorescence revealed that degradation promoted by **G-6599** was dependent on the ubiquitin proteasome pathway and the cullin-RING ligase (CRL) family, in addition to monovalent degrader binding, as indicated by the rescue of SMARCA2/A4 nuclear intensity using: inhibitors of the E1 activating enzyme (MLN7243), the proteasome (MG132), neddylation (MLN4924) and the parent SMARCA2/A4 bromodomain ligand (compound **7**) (Fig. 1c). Quantitative di-glycine (K-ε-GG) remnant profiling by mass spectrometry following treatment of cells with **G-6599** revealed cellular ubiquitination of multiple lysine residues on the expected target proteins, SMARCA2/A4 and PBRM1, with no off-target ubiquitination events (Fig 1d, Extended Data Fig. 2, Supplementary Table 2). The SWI/SNF PBAF protein, PBRM1, encodes a bromodomain with 46% homology to the SMARCA2/A4 bromodomain and is also an observed off-target of PROTACs of these proteins^27,29^. Similar to what has been observed with SMARCA2/A4 PROTACs, we observed **G-6599**-mediated ubiquitination of multiple lysines on both SMARCA2/A4, with the strongest ubiquitination occurring at C-terminal lysines mapping within or proximal to the bromodomain (Extended Data Fig 2). No ubiquitination of core SWI/SNF complex components was observed. An analysis of the global proteome further supported the selectivity, as SMARCA2/A4 and PBRM1 represented the only proteins depleted upon treatment of cells with **G-6599** (Fig 1e, Supplementary Table 3).

### G-6599 demonstrates similar cellular activity to potent reported bivalent degraders

We additionally compared the cellular activity of **G-6599** to previously reported VHL-based bivalent degraders targeting these proteins and/or FHD-286, a SMARCA2/A4 ATPase inhibitor currently in clinical development^27–30,32^. In SMARCA2/A4 -proficient SW1573 cells, **G-6599** exhibited comparable degradation potencies with the most potent PROTAC reported to date, A947 (Fig. 2a, Supplementary Table 4). The anti-proliferative effect of these molecules was tested in two disease model contexts, AR-dependent prostate cancer and SMARCA4-mutant non-small cell lung cancer (NSCLC). **G-6599** treatment resulted in a comparable inhibition of cell growth relative to A947 or FHD-286 in the SMARCA4-mutant models, NCI-H1944 and HCC515 (Fig. 2b, Supplementary Table 4). Although **G-6599** exhibited a slightly reduced potency on cell growth relative to A947 in AR-dependent prostate models (∼4-8 -fold shift in IC_50_), activity was comparable to FHD-286 (Fig. 2b, Supplementary Table 4). **G-6599** activity in the normal prostate epithelial line, RPWE-1, was significantly reduced relative to the AR+ prostate cancer lines, as was previously reported for a SWI/SNF inhibitor (Supplementary Table 4)^28^. Importantly, **G-6599** had no impact on cell growth in NCI-H522 cells that lack SMARCA2/A4^27^, further supporting the specificity of these molecules (Fig. 2b).

**Fig. 2.**
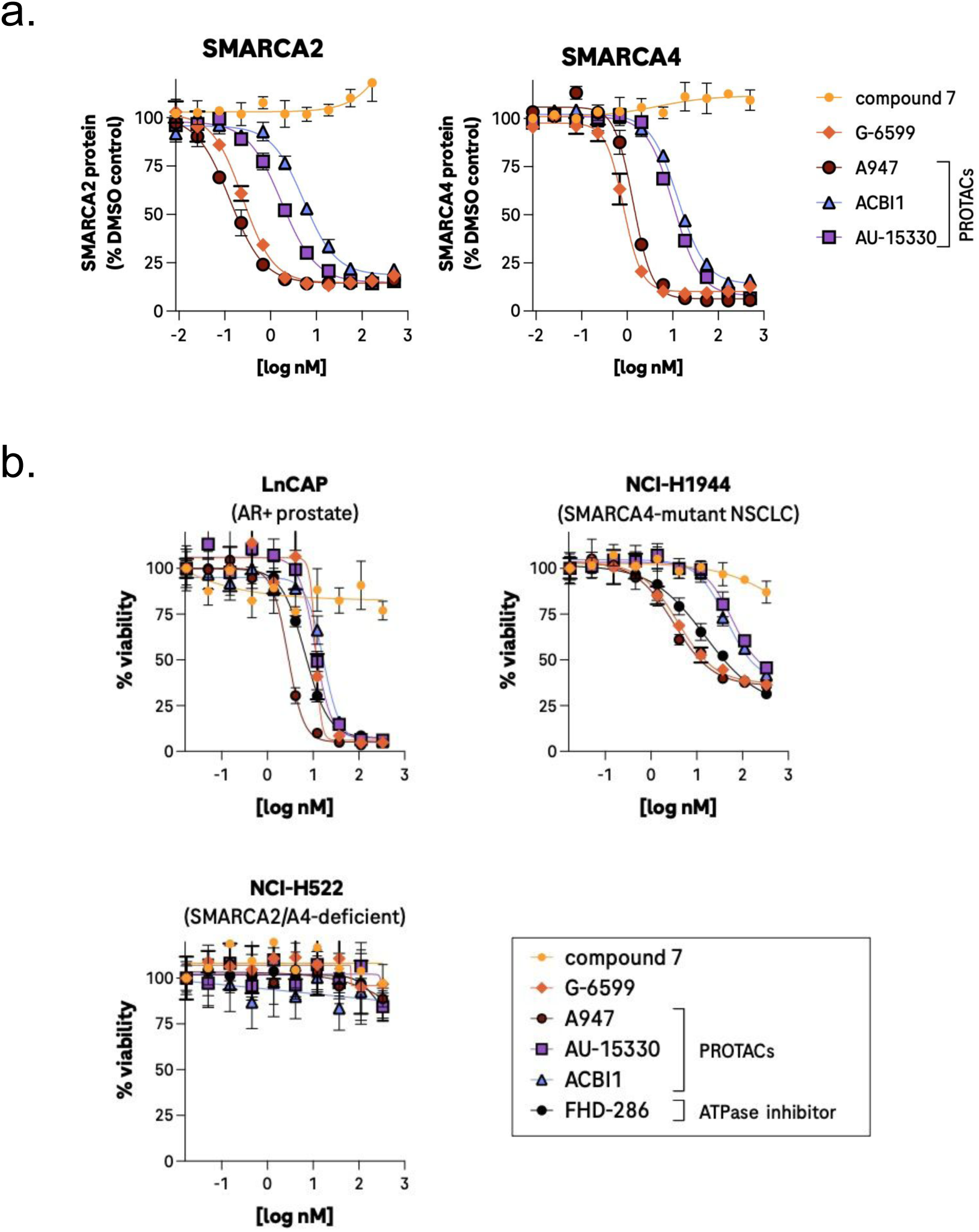
Monovalent degrader, G-6599, exhibits comparable potency and cellular activity with SMARCA2/A4 PROTACs and ATPase inhibitors. **a,** Dose-response curves comparing **G-6599** with bivalent SMARCA2/A4 PROTACs on SMARCA2 (left) and SMARCA4 (right) levels in SW1573 cells. Data are presented as mean ± s.d from triplicate cultures. **b,** Cellular viability following treatment with **G-6599**, PROTACs and the ATPase inhibitor FHD-286 in an AR-dependent prostate cell line (LnCAP), a SMARCA4-mutant NSCLC cell line (NCI-H1944) and in cells deficient for SMARCA2/A4 expression (NCI-H522). Data are presented as mean ± s.d from quadruplicate cultures.

### Monovalent degraders mediate protein depletion dependent on FBXO22 cullin-RING ligase

In order to elucidate the mechanism(s) of degradation, we carried out an array-based CRISPR screen with compound **4**, the best monovalent degrader available at the time, with the aim to identify genes required for compound **4** -mediated degradation of SMARCA2. As a control for the screen, we initially assessed which cullin may be required for compound **4** -mediated degradation by siRNA, given the requirement for neddylation (Extended Data Fig. 3a). Knockdown of Cullin-1 (CUL1) could partially overcome the effect of compound **4**, which not only enabled us to incorporate guide RNAs targeting CUL1 into our screen as a positive control, but indicated that degradation was likely occurring through a CUL1 associated substrate recognition subunit. NCI-H1944 cells stably expressing Cas9 were transfected in arrayed format with guide RNAs targeting 1384 genes involved in protein regulation, and levels of SMARCA2 were monitored by immunofluorescence in the presence of DMSO or compound **4**. As controls, guide RNAs targeting the olfactory receptor proteins, OR5M9 and OR6N1, had no impact on compound **4**-mediated degradation of SMARCA2, whereas guide RNAs targeting CUL1 could block compound **4**-mediated degradation (Fig. 3a, Supplementary Table 5). Guide RNAs that nonspecifically impacted SMARCA2 protein levels independent of drug treatment (ex. PSMD6) were excluded from the analysis. We identified five genes whose knockdown specifically prevented compound **4**-mediated degradation of SMARCA2. Three of the identified genes have been implicated in general regulation of CRL activation/function (UBA3, COPS5, CAND1), however, importantly, we determined that knockout of the CUL1 substrate adapter, S-phase kinase-associated protein 1 (SKP1), and the CUL1 substrate recognition subunit, FBXO22, were able to overcome compound **4**-mediated degradation (Fig. 3a). FBXO22 is one of approximately 70 F-box proteins and is the only F-box protein to encode a FIST domain (F-box and intracellular signal transduction proteins) that is present in many microbial sensory proteins (see schematic, Extended Data Fig. 3f)^35^. It is ubiquitously expressed across normal and malignant tissues (Extended Data Fig. 3b). To initially validate these findings, we utilized siRNAs targeting FBXO22 and SKP1 to demonstrate that knockdown could overcome compound **4**-mediated degradation of both SMARCA2 and SMARCA4 in a second cell line, SW1573 (Fig. 3b). siRNA mediated knockdown of FBXO22 could reverse compound **4**-mediated degradation of SMARCA2 at all doses tested, whereas the siRNA-mediated knockdown of a control CUL1 CRL, FBXO42, had no impact on monovalent degrader activity (Extended Data Fig. 3c, d). Importantly, genetic knockout of FBXO22 was able to largely abrogate activity of **G-6599** in SMARCA4-mutant HCC515 cells, however had no impact on degradation mediated by the VHL-based PROTAC A947 (Fig. 3c, Extended Data Fig. 3e). Re-expression of wild-type (WT) FBXO22 in FBXO22-knockout HCC515 cells was able to restore **G-6599**-mediated degradation, with no influence on activity of the PROTAC A947 (Fig. 3d, Extended Data Fig. 3f-h). However, an FBXO22 mutant that cannot associate with SKP1 (FBXO22-ΔFbox) was unable to restore monovalent degrader function, demonstrating the requirement for FBXO22 ligase activity in monovalent degrader function. To address whether recognition of full-length SMARCA2/A4 protein is necessary for degradation, we expressed the bromodomain (BD) of SMARCA2 (aa. boundaries, S1377-Q1486; HA-tag) in cells (Extended Data Fig. 3i). Treatment of cells with **G-6599** resulted in the dose-dependent degradation of the HA-tagged SMARCA2-BD that was comparable to degradation observed with the PROTAC (Fig. 3e). Degradation of the HA-tagged bromodomain was additionally neddylation-dependent (Extended Data Fig. 3j). Overall, these results indicate that the FBXO22/SKP1/CUL1 ligase is required for SMARCA2/A4 monovalent degrader activity.

**Fig. 3.**
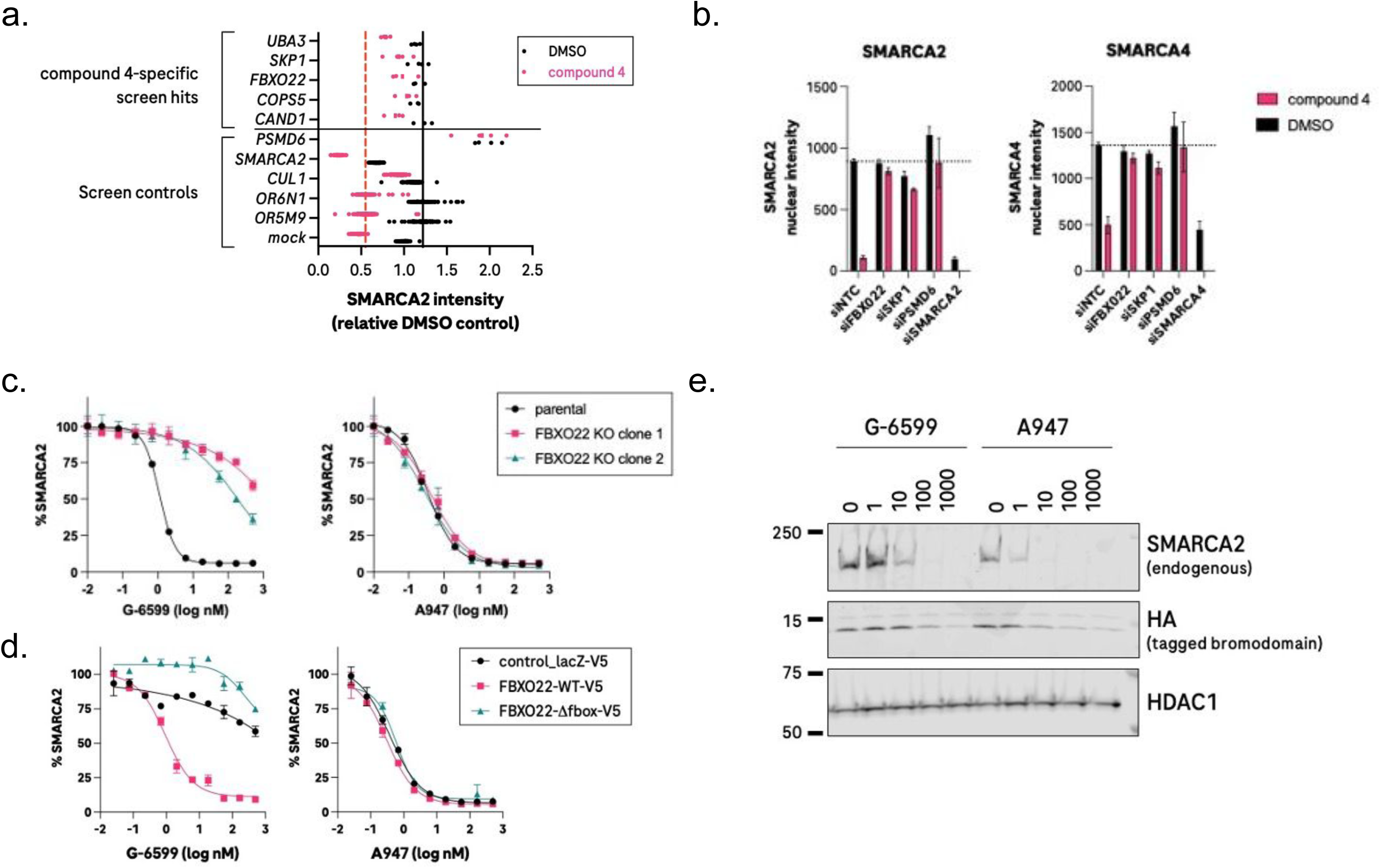
FBXO22 ligase activity is required for SMARCA2/A4 degradation. **a,** Arrayed CRISPR screen to identify modifiers of compound 4 -mediated degradation of SMARCA2 by immunofluorescence in NCI-H1944 cells stably expressing Cas9. The effect of guide RNAs targeting the respective gene on SMARCA2 expression in the presence of DMSO control and compound 4 (500 nM). Data points reflect individual replicate cultures (3-4 replicates for test conditions and 76-80 replicates for plate controls), with the lines representing the median intensity levels for the OR6N1 and OR5M9 gRNA controls in the DMSO (black) or compound 4 (red) conditions. Nuclear intensity of SMARCA2 is presented relative to DMSO control conditions. **b,** Immunofluorescent-based detection of SMARCA2 and SMARCA4 in SW1573 cells transfected with the respective siRNAs. Cells were treated for 8 h with 200 nM compound **4** or DMSO control. **c,** Effect of varying concentrations of **G-6599** or PROTAC control (A947) on SMARCA2 levels in isogenic parental and FBXO22 knockout HCC515 cells. **d,** Effect of varying concentrations of **G-6599** or PROTAC control, A947, on FBXO22 knockout HCC515 cells reconstituted with V5-tagged wild-type or ligase-dead (ΔFbox) FBXO22. Nuclear intensity of SMARCA2 within V5-positive cells is presented relative to DMSO control conditions. **e,** Degradation of the HA-tagged SMARCA2 bromodomain expressed in HCC515 cells following overnight treatment with varying concentrations of the respective molecules.

### G-6599 induces a ternary complex between SMARCA2 and FBXO22

To address whether SMARCA2/A4 monovalent degraders result in ternary complex formation in cells, we initially utilized a proximity labeling approach whereby the C-terminus of SMARCA2 was tagged with the biotin ligase “TurboID” and expressed in cells in a doxycycline(dox)-inducible manner along with V5-tagged FBXO22 (Fig. 4a). In the presence of MLN4924, added to prevent neddylation and degradation of SMARCA2-TurboID, **G-6599** treatment resulted in the biotinylation of FBXO22, with no detectable biotinylation in the absence of the monovalent degrader (Fig. 4b). **G-6599**-mediated biotinylation was additionally observed in cells expressing FBXO22-ΔFbox, and biotinylation could be outcompeted by treatment of cells with molar excess of the parent ligand (compound **7**) (Extended Data Fig. 4a, b). We additionally assessed ternary complex formation via NanoBRET^®^ in FBXO22 knockout HCC515 cells expressing SMARCA2-nanoLuc and FBXO22-HaloTag. **G-6599** treatment resulted in a dose-dependent increase in the BRET signal that was not observed with bivalent degrader A947 treatment (Fig. 4c). Finally, we also studied target engagement in cell lysates by a cellular thermal shift assay. Addition of **G-6599** to lysates of HCC515 expressing a V5-tagged FBXO22 resulted in a change in the thermal stability of FBXO22 that was not observed with addition of the parental SMARCA2/A4 ligand (compound **7**) (Fig. 4 d,e). Taken together, the data supports target engagement and ternary complex formation of FBXO22 and SMARCA2 in cells mediated by monovalent degrader **G-6599**.

**Fig. 4.**
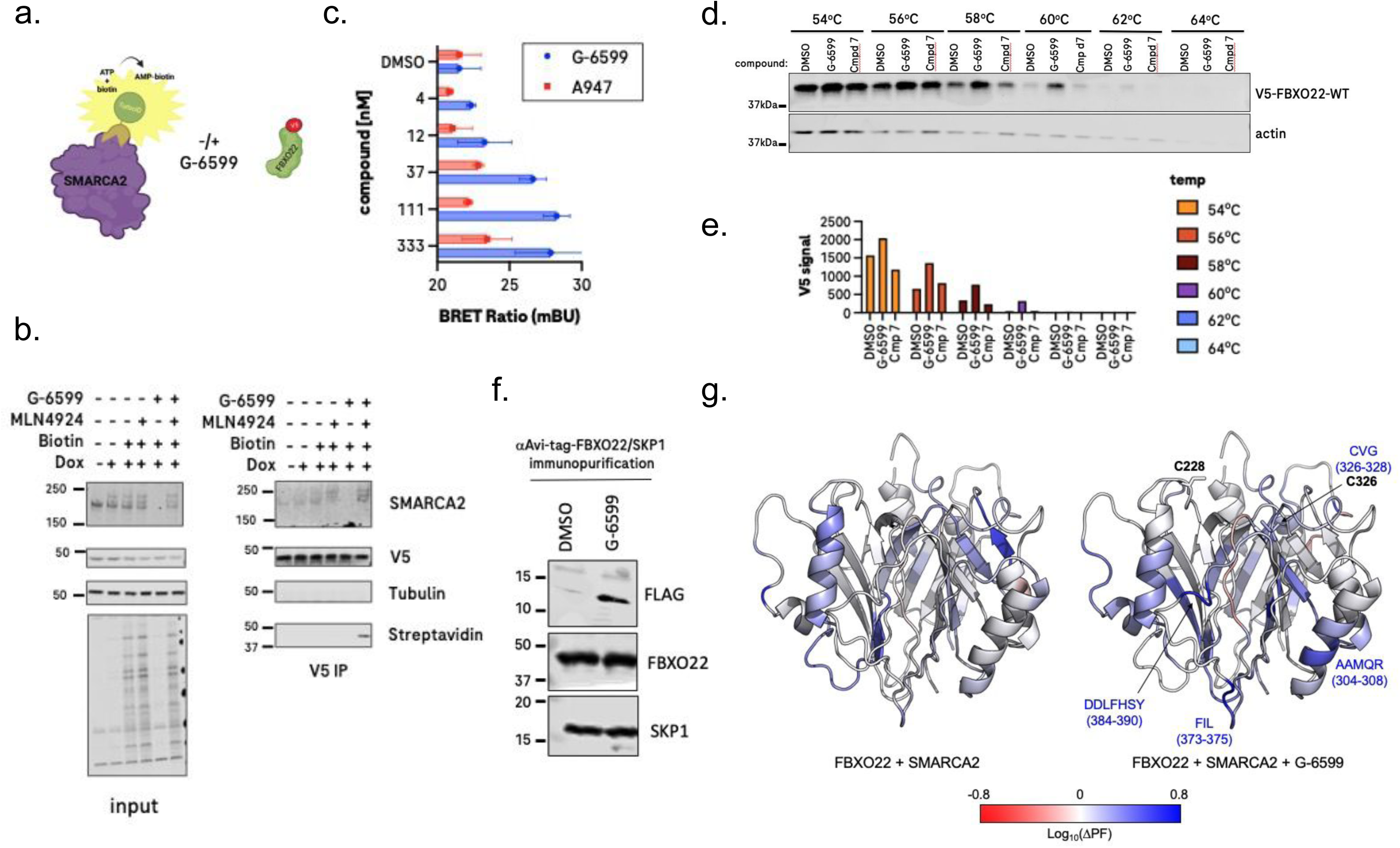
SMARCA2 monovalent degraders recruit FBXO22. **a,** Depiction of proximity-labeling approach to evaluate cellular engagement of FBXO22 with SMARCA2. **b**, FBXO22 knockout HCC515 cells stably expressing V5-tagged FBXO22 and doxycycline-inducible SMARCA2-TurboID were pre-treated for 72 h with 1 µg/mL doxycycline followed by a 1 h treatment with 6 µM MLN4924 and subsequent addition of 200 nM **G-6599** and 50 µM biotin overnight. Biotinylated proteins were detected by streptavidin Western blotting of input and anti-V5 immunoprecipitates. Tubulin served as a negative control protein in V5 immunoprecipitates. **c,** BRET ratios in FBXO22 knockout HCC515 cells transiently transfected with SMARCA2-nanoLuc and FBXO22-HaloTag following overnight treatment with DMSO or varying concentrations of of **G-6599** and control PROTAC, A947. Cells were pretreated for 1 h with 10 µM MLN4924 and maintained in MLN4924 for the duration of the experiment. **d**, Representative anti-V5 immunoblot from lysates from FBXO22 knockout HCC515 cells reconstituted with V5-tagged FBXO22 treated with the respective compounds after exposure to a 54 °C-64 °C temperature range. Actin served as a loading control. **e**, Licor-based quantification of V5 immunoblots from 4e **f,** Monovalent degrader-mediated co-immunoprecipitation of recombinant FBXO22 protein with recombinant SMARCA2 bromodomain in solution. Recombinant Avi-tagged FBXO22/SKP1 protein immobilized on anti-Avi resin was mixed for 1 h with FLAG-tagged SMARCA2 protein (aa. S1377-Q1486) in DMSO or 1 µM monovalent degrader. Precipitates were washed and immunoblotted for the respective proteins. **g**, Hydrogen-deuterium exchange protection factor change (Log_10_ΔPF) upon treatment of recombinant SMARCA2 bromodomain with FBXO22 (left) and in the presence of **G-6599** (right) compared FBXO22 alone. Colored map overlaid on FBXO22 cryo-EM structure (PDB: 8UA3, BACH1 BTB omitted for clarity). Red = reduced protection / increased deuterium uptake, blue = increased protection / reduced deuterium uptake, white = no change. Residues for which a significant increase in protection from deuterium uptake in the presence of **G-6599** are indicated in blue (right). Location of cysteines 228 and 326 is indicated (right).

To further address the question of a potential physical interaction mediated by SMARCA2/A4 monovalent degraders, we assessed whether FBXO22 can interact with SMARCA2 in solution through affinity purification. Avi-tagged or FLAG-tagged FBXO22 was co-produced with SKP1 and utilized as a bait protein in pulldowns with FLAG-tagged or His-tagged SMARCA2 bromodomain, respectively (Extended Data Fig. 4 d, e). Epitope-tagged SMARCA2 was enriched in FBXO22 pulldowns in the presence of **G-6599** relative to DMSO (Fig. 4f and Extended Data Fig. 4f, g). To reach a better understanding of the conformational dynamics of the ternary complex and the regions of FBXO22 involved in protein-protein contacts with SMARCA2 in the presence of **G-6599**, we carried out hydrogen-deuterium exchange mass spectrometry (HDX-MS) experiments. The extent of the protection from hydrogen-deuterium exchange near the C-terminal region of the FIST domain of FBXO22 in the presence of SMARCA2 and **G-6599** compared to the absence of compound suggests that this region of FBXO22 may be involved in the ternary complex (Fig. 4g, Extended data Fig. 4h, Supplementary Table 6). Some of the areas where we observed less deuterium intake in the presence of **G-6599**, such as Ala304-Arg308, Phe373-Leu375 and Asp384-Tyr390, are part of the FBXO22 FIST-3 domain (or FIST-C domain) that was described to interact with endogenous substrates, such as BACH1^36,37^. In addition, we found significant protection from deuterium incorporation near Cys326 (Cys326-Gly328), a cysteine residue that was targeted by reported covalent monovalent degraders of NSD2 and FKBP12 acting through FBXO22^38,39^.

### G-6599 acts as a monovalent degrader by covalently targeting FBXO22

Considering the involvement of the Cys326 region in covalent monovalent degraders recruiting FBXO22 and that 2-alkynylpyridines have been reported as covalent warheads modifying cysteine residues^40,41^, we sought to establish if our series of monovalent degraders, including **G-6599**, was operating via a similar mode of action. We observed a moderate correlation between cysteine reactivity *in vitro* and degradation potency for monovalent degraders containing an alkyne, supporting a potential covalent mechanism (Fig. 5a). Interestingly, **G-6599** was an outlier, achieving more potent degradation while being less reactive toward cysteine *in vitro*. We hypothesized that the azepane tail may promote additional hydrophobic contacts with the FBXO22 protein surface, diminishing reliance on intrinsic cysteine reactivity to form a strong ternary complex to achieve potent SMARCA2 degradation. To determine if a covalent adduct was formed, we conducted an intact protein mass spectrometry experiment using recombinant SKP1-FBXO22 in the presence and absence of SMARCA2 bromodomain (Fig. 5b). We observed SMARCA2-independent formation of a single and, to a lesser extent, a double adduct of **G-6599** in solution. The lack of dependency on the presence of SMARCA2 to promote a covalent adduct may be explained by the high protein and compound concentrations under the experimental conditions, however, we cannot exclude the possibility that the monovalent degrader tail may have affinity on its own for FBXO22. Peptide mapping in the presence of SMARCA2 revealed the modification of three cysteine residues: Cys326, Cys228 and Cys117 (Fig. 5c, Extended data Fig. 5a, Supplementary Table 7). Cys228 (and Cys227) has also been reported in the context of targeted protein degradation of FKBP12 via FBXO22 using a bivalent degrader approach^42^. Cys117 is located in a solvent-exposed loop where no significant protection from hydrogen-deuterium exchange was observed and is thought to be an unspecific modification under the *in vitro* assay conditions (Fig. 4g, Extended data Fig. 4h). Contrary to recent reports of monovalent degraders acting through the recruitment of Cys326 of FBXO22 via an aldehyde formed by monoamine oxidase biotransformation of a primary amine in cells^38,39^, our data do not support that **G-6599** is operating via a similar mechanism. Differently from monovalent degrader **5**, possessing a primary amine moiety similar to these reported monovalent degraders, no aldehyde was formed for **G-6599** after treatment for 4 hours in cell media containing 10% of fetal bovine serum (FBS) or upon treatment in HCC515 cells (Extended data Fig. 5b). In addition, absence of FBS in cell media or co-treatment with monoamine oxidase inhibitors did not rescue degradation promoted by **G-6599** or the VHL based bivalent degrader control A947. Conversely, the above conditions abrogated SMARCA2 degradation with compound **5**, presumably through preventing the transformation to an aldehyde (Extended data Fig. 5 c, d). Instead, **G-6599** likely forms a covalent adduct with FBXO22 cysteine residue(s) through its electron-deficient 2-alkynylpyridine moiety (Fig. 5b). Intrigued by these findings, we next sought to determine if these cysteine residues were involved in the degradation of our monovalent degrader. FBXO22-knockout HCC515 cells were engineered to re-express V5-tagged wild-type or mutant FBXO22, and **G-6599**-mediated degradation was monitored in cells expressing comparable levels of V5 (Figure 5d, Extended data Fig 5 e, f). Modification of Cys326 or Cys228 to alanine resulted in a significant reduction in degradation mediated by **G-6599**, whereas modification of Cys227 to alanine had minimal impact on **G-6599** -mediated degradation of SMARCA2. The levels of **G-6599** -mediated degradation of SMARCA2 were comparable between cells expressing the Cys228Ala mutant relative to the double mutation of Cys227Ala/Cys228Ala, further indicating a lack of involvement of Cys227. SMARCA2 degradation mediated by the control VHL PROTAC, A947, was unaffected by wild-type or mutant FBXO22 (Extended data Fig. 5g). Consistent with these degradation findings, we observed no thermal cellular stability shifts mediated by **G-6599** in lysates from cells expressing FBXO22 Cys228Ala or Cys326Ala, compared to FBXO22 wild-type expressing cells (Fig. 5e vs Fig 4 d, e). While quantification of the peptide sequence modification of FBXO22 by mass spectrometry in the presence of SMARCA2 indicates that Cys228 is the primary site of modification over Cys326 (Supplementary Table 7; % modification: 28.22% and 0.09%, respectively, using trypsin digestion), the cysteine mutant data and the HDX-MS experiments showing greater protection from hydrogen-deuterium exchange near Cys228 or directly in the region where Cys326 is located (Fig. 4g right, Fig. 5d) suggest that both cysteines are critical for efficient turnover of SMARCA2 promoted by the monovalent degrader. The modification of Cys326 observed in the peptide mapping experiment may not be biologically relevant and can possibly be explained by the high concentrations used in the biochemical experiment. However, we cannot exclude with certainty a mechanism that involves some level of covalent modification of Cys326. Our data, in addition to reports from other groups^38,39,42^, showing that multiple cysteines located on FBXO22 are prone to covalent modification leading to favorable protein-protein interactions and degradation of the recruited target, support the notion that FBXO22 possesses a “high degree of superficial ligandability”^43^.

**Fig. 5.**
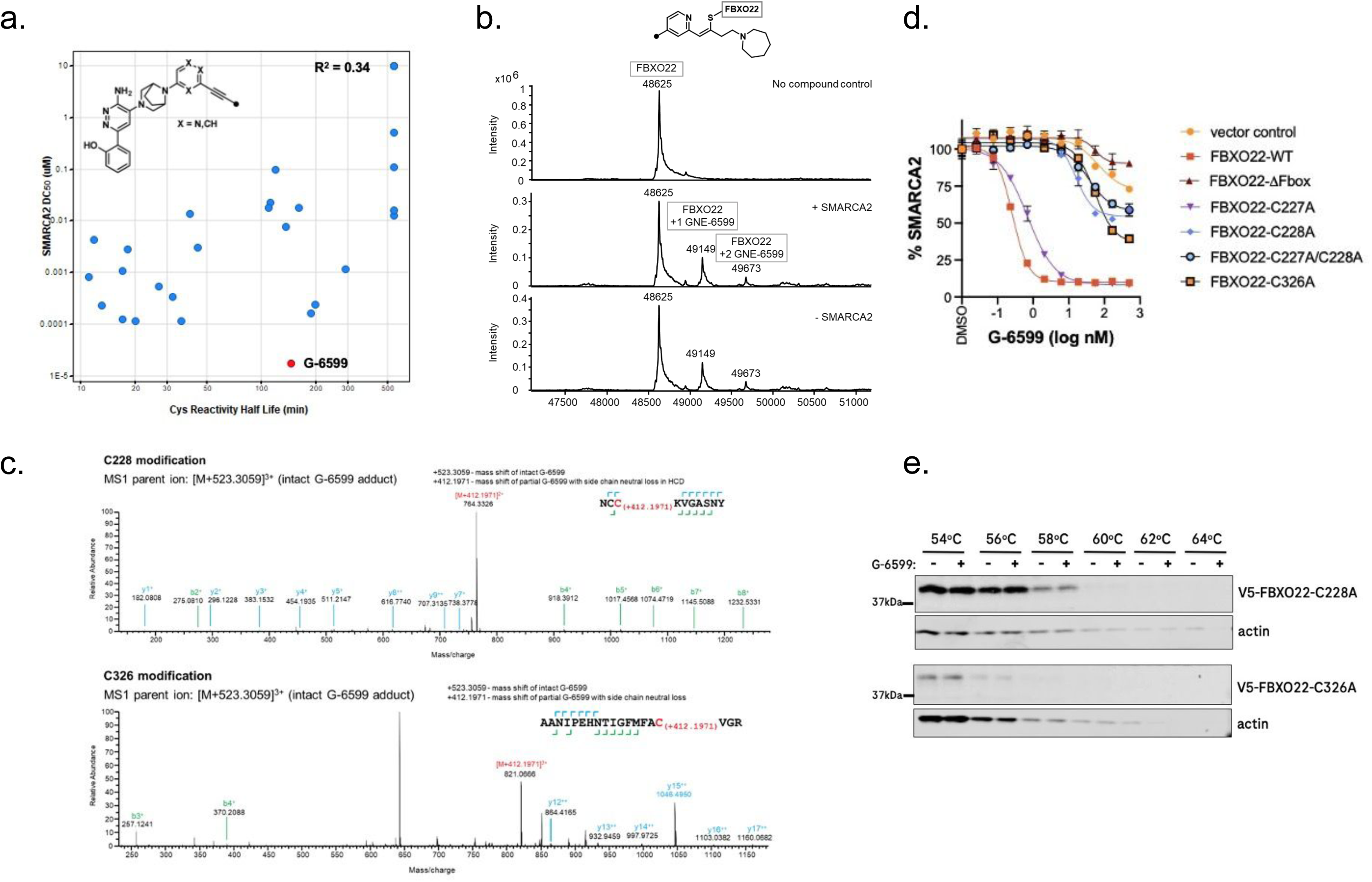
The recruitment of FBXO22 by G-6599 involves FBXO22 cysteine modification. **a,** Correlation between SMARCA2 degradation potency (DC_50_) and *in vitro* cysteine reactivity half life for a series heteroaryl alkyne monovalent degraders. **b,** Intact mass spectrometry of 1 µM of FBXO22/SKP1 alone (top) or in the presence of 20 µM of **G-6599** with (middle) or without (bottom) 1 µM of SMARCA2 bromodomain. **c,** Identification of covalently modified FBXO22 peptides using mass spectrometry after 4 h treatment of 1 µM FBXO22/SKP1 and SMARCA2 bromodomain in the presence of 50 µM of **G-6599** for Cys228 (top) and Cys326 (bottom) modification. **d,** Effect of varying concentrations of **G-6599** on SMARCA2 levels in FBXO22 knockout HCC515 cells reconstituted with V5-tagged wild-type or mutated forms of FBXO22. Nuclear intensity of SMARCA2 within V5-positive cells is presented relative to DMSO control conditions. **e,** anti-V5 immunoblot from lysates of HCC515 cells expressing V5-tagged FBXO22 mutants treated with **G-6599** or DMSO control after exposure to a 54-64 °C temperature range. Actin served as a loading control.

## Discussion

Herein, we report the discovery and optimization of monovalent degraders of SMARCA2/A4 via a deliberate chemistry approach from a known functionally inactive bromodomain ligand. Parallel synthetic chemistry was leveraged to generate a targeted, but diverse, screening library that was evaluated in a high-throughput degradation assay and led to the identification of hits with weak degradation activity. Further chemical diversification demonstrated that it is possible to improve degradation potency of such starting points via traditional structure-activity-relationship exploration. **G-6599** emerged as an exceptionally potent and proteome-wide selective monovalent degrader of SMARCA2/A4. The degradation potency (DC_50_) of the original screening hits, compounds **2** and **3**, was improved by 1000- and 100-fold for SMARCA2 and SMARCA4, respectively, compared to **G-6599**. **G-6599** demonstrated similar degradation activity to reported VHL-based bivalent degraders, in addition to comparable and specific anti-proliferative activity in relevant models relative to the bivalent degraders and the ATPase inhibitor FHD-286.

Investigation of the mode of action revealed that **G-6599** is operating through the ubiquitin proteasome system and a cullin-RING E3 ligase. A CRISPR screen using an earlier generation analog, compound **4**, suggested that the cullin-RING E3 ligase complex FBXO22 is involved in the degradation by this class of molecules. FBXO22 is a broadly expressed E3 ligase across normal and malignant tissues, and as such, would be a relevant E3 ligase to harness in the context of oncology. We later confirmed that degradation promoted by our most potent monovalent degrader **G-6599** was also dependent on FBXO22 using mutant KO cell lines. We were able to validate the direct recruitment of FBXO22 to SMARCA2 in the presence of **G-6599** using biochemical methods and cellular experiments, including co-immunoprecipitation, TurboID, NanoBRET^®^ and thermal shift. Interrogation of the FBXO22 protein surface by HDX-MS suggested that the presence of the monovalent degrader bound to SMARCA2 promotes a distinct and more sustained interaction compared to SMARCA2 alone. The interface involved in protein-protein contacts promoted by **G-6599** is located in the FIST-3 domain of FBXO22, a region that is also involved in the binding of the endogenous substrate BACH1. While we observed increased protection from deuterium uptake in the presence of SMARCA2 without **G-6599**, suggesting a potential weak interaction, we were not able to demonstrate that SMARCA2 has basal affinity for FBXO22 using TurboID and BLI (data not shown). We cannot exclude that this increase in protection in the presence of SMARCA2 alone may be caused by the high concentrations used in the biochemical experiment and may not be relevant in the cellular context.

Motivated by recent reports of mono- and bivalent protein degraders covalently recruiting FBXO22 and by an emerging correlation between degradation potency and cysteine reactivity *in vitro* with our monovalent degraders, we investigated this mode of action further. Using intact mass spectrometry, we could confirm formation of FBXO22 covalent adducts in the presence of **G-6599**. Interestingly, the formation of the covalent adducts was independent of the presence of SMARCA2. We cannot rule out the possibility that **G-6599** may have intrinsic binding affinity to FBXO22 alone, however, we believe that this is likely unspecific and caused by the high concentration used in the experiment. Peptide mapping revealed the modification of three cysteine residues on FBXO22: Cys117, Cys228 and Cys326. Cys117 was located in a highly solvent-exposed loop, and modification is thought to be unspecific under the assay conditions. While Cys228 was identified as the primary site of conjugation, HDX-MS suggests that the region containing Cys326 is more directly involved in protein-protein interactions. Notably, several groups independently reported both cysteines to be involved in the FBXO22-mediated degradation of different proteins. Monovalent degraders of NSD2, FKBP12 and XIAP possessing a primary amine serving as an aldehyde pro-drug via biotransformation were reported to lead to the recruitment of FBXO22 Cys326. Conversely, Cys228 was reported to be involved in the degradation of FKBP12 using a heterobifunctional molecule containing a FBXO22 binding ligand bearing a chloroacetamide warhead. In the latter example, degradation was dependent on Cys228, but also on the adjacent Cys227. In our case, we did not observe significant dependency on Cys227. However, we found that both single mutants C228A and C326A can lead to significant, but incomplete rescue of degradation, suggesting that both cysteines are important to achieve recruitment of FBXO22 to SMARCA2 for degradation upon treatment with **G-6599**. However, we believe that Cys228 may play a direct role in engaging **G-6599** covalently, while Cys326 may be more important in creating a favorable protein surface for interacting with SMARCA2 without interacting covalently with the small molecule degrader. In addition, we demonstrated that **G-6599** does not undergo biotransformation to covalently promote a ternary complex with FBXO22, but is likely to react via its electron-deficient alkyne. This demonstrates that **G-6599** is a highly potent and specific monovalent degrader of SMARCA2/A4 possessing a unique mechanism compared to other reported degraders operating through FBXO22, but also that FBXO22 has a high level of processivity and potentially a broad substrate scope.

Recent developments in the discovery of monovalent degraders from established protein ligands offer great promise in the development of new therapies. They offer the opportunity to combine the benefits of protein degradation - i.e.: longer duration of action, potential for targeting scaffolding function of proteins and increased selectivity - with the more favorable physicochemical and pharmacokinetic properties of small-molecule drugs. The discovery of monovalent degraders also offers the opportunity to find previously unexploited degradation machinery and expand the repertoire of E3 ligases co-optable for targeted protein degradation. Moreover, taking advantage of the covalent mode of action on the E3 ligase may provide additional benefit by supercharging the E3 ligase with the monovalent degrader leading to the formation of a pseudo-binary complex with the target protein, increasing degradation efficiency.

While challenges remain to realize the full potential of this novel approach, particularly in addressing the currently empirical discovery process, we believe that the therapeutic potential of target-anchored monovalent degraders warrants continued exploration.

## Methods

### Cell Lines

All cell lines were grown in RPMI 1640 supplemented with 10% fetal bovine serum (FBS), 2 mM L-Glutamine and 100 U/mL penicillin-streptomycin (Gibco) under 5% CO_2_ at 37°C. Cell lines were obtained from American Type Culture Collection (ATCC) (NCI-H1944, SW1573, LNCAP, and NCI-H522) or licensed from UT Southwestern Medical Center (UTSW) (HCC515). Cell line identity was verified by high-throughput single nucleotide polymorphism (SNP) genotyping using Illumina Golden Gate multiplexed assays. SNP profiles were compared to SNP calls from internal and external databases to determine or confirm ancestry. All cell lines tested negative for mycoplasma contamination prior to storage/use at our institute.

### Engineered Cell Lines

FBXO22 knockout HCC515 knockout lines were generated by CRISPR/Cas9 through lentiviral delivery of a guideRNA targeting FBXO22 (gFBXO22), TTCGTGTTGAGTAACCTGG. Cells expressing gFBXO22 were selected with puromycin for 7 days. When puromycin resistance was acquired, cells were transfected with TrueCut Cas9 v2 (Thermo Fisher Scientific) using Lipofectamine CRISPMAX (Thermo Fisher Scientific) for 48 h and clones were subsequently established by serial dilution and screened for FBXO22 ablation by Western.

The pLenti6.3 system was utilized for ectopic expression of wild-type/mutant FBXO22 in FBXO22-knockout HCC515 cells, as well as ectopic expression of the SMARCA2 bromodomain (S1377-Q1486) in parental HCC515 cells. All FBXO22 DNA constructs were generated with a C-terminal V5 tag and the SMARCA2 bromodomain was engineered with a C-terminal HA tag. The pMinDucer system was utilized for ectopic expression of a SMARCA2 construct engineered with a C’terminal TurboID tag.

### Antibodies

The following antibodies were utilized: SMARCA2 (Cell Signaling, 11966, dilution 1:2000), SMARCA4 (Abcam, ab110641, dilution 1:1000), HA (Cell Signaling, 2367, dilution 1/1000), HDAC1 (Cell Signaling, 34589, dilution 1:1000), V5 (Cell Signaling, 80076, dilution 1/1000), V5 (Chromoteck, v5tma-20, dilution 1/1000), Tubulin (Cell Signaling, 2128, dilution 1:1000), FLAG (Cell Signaling, 14793, dilution 1/1000), FLAG (Sigma-Aldrich, M8823, dilution 1:1000), FBXO22 (Abcam, ab230395, dilution 1/1000), SKP1 (Cell Signaling, 12248, dilution 1/1000), CUL1 (Invitrogen, 71-8700, dilution 1/1000), and His (Cell Signaling, 12698, dilution 1/1000)

### Reagents

The chemical compounds MLN7243, MLN4924 and MG132 were obtained from SelleckChem. The chemical compounds ACBI, AU-15330, FHD-286 and Marplan were synthesized in-house according to the published procedures. PXS-4728A was acquired from MedChemExpress (HY-112726). Aminoguanidine was purchased from Millipore (396494). Biotin (230095000) and Optimem (11058021) were both purchased from Thermo Fisher.

### Cell Viability

Cells were plated in 384-well plates at 1000-1500 cells per well and treated the next day with the indicated concentration of compounds. Following 7 days of treatment, viability was determined by using CellTiter-Glo Luminescent cell viability assay (Promega, G7573). CellTiter-Glo solution (30 µl) was added directly to the cells, mixed and incubated for 10 min. Luminescence was measured using the multimode plate reader EnVision 2105 (PerkinElmer). Viability was normalized to DMSO treated control cells. Cell viability experiments were performed in quadruplicate cultures.

### Immunofluorescence

Cells (3000) were plated in 384-well plates (PhenoPlate™ 384-well microplates, PerkinElmer) overnight. Cells were subsequently treated for 20 h with the indicated concentration of compounds. Cells were fixed with 4% formaldehyde for 15 min at room temperature and plates were washed three times with PBS and incubated with a blocking solution (10% FCS, 1% BSA, 0.1% TritonX-100, 0.01% Azide in PBS) for 1.5 h. Following PBS wash (3x), primary antibody diluted 1:1000 in blocking buffer was added and incubated overnight at 4°C. Following 3 times washing in PBS, plates were treated with secondary antibodies (rabbit-Alexa 488, Thermo Fisher A21206, 1:1000) for 1 h at room temperature in the dark. Plates were washed 3 times in PBS. Hoechst 33258 (Thermo Fisher H3569) at 1:5000 was added to the wells and the plates were incubated for 10 min and imaged on an Opera Phenix™ High Content Screening System (PerkinElmer). Using Hoechst H3569 nuclear staining as a mask, nuclear SMARCA2 and SMARCA4 mean signal intensity were quantified. No primary antibody served as a background signal control. Immunofluorescence experiments were performed in triplicate cultures.

### High-throughput immunofluorescence assay

SW1573 cell lines were used for compound treatment. Cell lines were cultured in Complete media (RPMI1640 + 10% Tet-Free HI FBS + 1X L-Glutamine + 1X Pen Strep. Compound libraries were prepared in 384 well LDV Echo plates as 10 mM in DMSO. Compounds were dispensed onto black PDL-treated CellCarrier-384 Ultra Microplates (Perkin Elmer 6057500) in a dose-response manner (0.1%. DMSO final) via Echo.

Cells were seeded at 2500 cells per well in 25 µL media into 384-well plates (Perkin Elmer) containing compounds using the ViaFlo robot. Cells were allowed to settle for 20 min at room temperature and then incubated for 18 h at 37 °C in a 5% CO_2_ humidified incubator. After compound treatment, cells were fixed with 4% PFA for 20 min at room temperature. Then, they were permeabilized and blocked with IF blocking solution (10% FCS, 1% BSA, 0.2% TritonX-100 in PBS, 0.01% Azide) at room temperature for 1 h. Primary antibodies SMARCA2 (Cell signaling Cat# 11966, 1:1200) and SMARCA4 (Rabbit monoclonal [EPNCIR111A] to SMARCA4 Abcam ab 110641) diluted in IF blocking buffer were added to the cells then cells were incubated overnight at 4 °C. The following day, cells were co-stained with a secondary antibody (Donkey anti-Rabbit Alexa Fluor 488 #A21206; 1:1000) and Hoechst 33258 (#H3569; 1:10000) diluted in PBS and incubated 1 h at room temperature in the dark. Images of the stained cells were acquired with a Cellomics CX7 or Opera Phenix Microscope. Total fluorescence intensity in nuclei in the green channel was measured and quantified. Further analysis and dose-response curves were fitted to obtain IC_50_ values, D_max_, and Z’ factors using Genedatta or GraphPad PRISM.

### RNAi

Cells (10,000/well) were reverse transfected using Lipofectamine RNAimax (Thermo Fisher Scientific) with 20 µM of siRNA. All siRNA were SMARTPOOL ON-TARGETPLUS siRNA (Thermo Fisher Scientific).

### Immunoblotting

Cells were lysed in RIPA buffer containing 1M NaCl and subsequently homogenized for 3 min at speed 10 (NextAdavance, Bullet BlenderR 24). Proteins (12 µg or 18 µg) were resolved on 4-12% Bis-Tris or 3-8% Tris-Acetate gels and transferred to nitrocellulose membranes by iBlot (Thermo Fisher Scientific). Membranes were incubated overnight at 4 °C in PBS with 5% Milk with the respective primary antibodies. IRDyeR (LI-COR) conjugated secondary antibodies were used to bind primary antibodies. Western blots were visualized on the Odyssey CLx Imager (LI-COR).

### Quantitative RT-PCR

RNA was isolated 48 h after transfection using the RNeasy Plus mini kits (Qiagen). Gene expression levels were determined by Taqman using the Taqman One-Step RT-PCR Master Mix Reagents kit (Thermo Fisher Scientific). The following Taqman gene expression assays were utilized: FBXO42 (Hs00398587_m1) and GUSB (Hs00939627_m1). Analysis was performed using 7900HT SDS (Thermo Fisher Scientific) and expression levels are presented relative to the housekeeping gene, GUSB (2^-ΔCt^).

### Recombinant protein expression and purification

Genes for FBXO22 and SKP1 were synthesized at Genscript with codon optimization for insect cell expression, and then subcloned into appropriate vector backbones. Recombinant baculoviruses were produced and amplified according to manufacturer’s protocols. Full length FLAG-tagged FBXO22-SKP1 complex was produced by infecting Sf9 insect cells with separate recombinant baculoviruses encoding tagged FBXO22 and untagged SKP1. Biotinylated full length His-Avi-FBXO22-SKP1 was produced by coinfecting His-Avi-FBXO22, untagged SKP1, and untagged BirA for in cell biotinylation. Cells were harvested 72 hours post infection and frozen until purification.

His-Avi-FBXO22-SKP1 cells were resuspended in lysis buffer (40 mM HEPES pH 7.5, 350 mM NaCl, 10% glycerol) and lysed by sonication. Lysate was clarified by ultracentrifugation, and the supernatant was then loaded onto a Ni-NTA column. Beads were washed with 10 column volumes (CVs) of lysis buffer supplemented with 40 mM imidazole. Bound proteins were eluted from the column with 5CVs of lysis buffer + 300 mM imidazole. Elution was concentrated and then applied to a Superdex 200 Increase 10/300 (Cytiva) for further purification. Fractions were run on SDS-PAGE, and visualized by Coomassie blue staining. Successful biotinylation of the Avi-tag was assessed by binding free streptavidin to His-Avi-FBXO22-SKP1 and checking for a supershift of the His-Avi-FBXO22 protein on SDS-PAGE. Biotinylation was ∼100%. Pure fractions were pooled and concentrated to ∼4 mg/mL, and then frozen in LN2 for future use.

FLAG-FBXO22-SKP1 was purified in a similar fashion, with the exception of the affinity purification step using M2 anti-FLAG agarose (Sigma). Proteins were eluted from the column using lysis buffer supplemented with FLAG peptide.

### Recombinant protein pull-down studies

Avi binding beads (made in-house) and Anti-FLAG M2 (Sigma-Aldrich, M8823) affinity resins were used for Avi and Flag pulldown, respectively^44^. Agarose beads for binding studies were pre-washed with binding buffer (20 mM HEPES pH 7.5, 200 mM NaCl, 5% glycerol). Following washes, 10 ug of recombinant Avi-tagged FBXO22/SKP1 or Flag-tagged FBXO22/SKP1 protein were immobilized on binding beads for 1 h at 4 °C with gentle rocking. Following immobilization, beads were washed (3x) in 300 µL binding buffer to remove any excess unbound protein. These beads were used for all further incubations with bait proteins and no bait controls. 40 µg of FLAG-tagged SMARCA2 or His-tagged SMARCA2 protein in the presence or absence of monovalent degrader (1 µM), PROTAC control (1 µM) or parent ligand (50 µM) were incubated with beads on ice in a total volume of no more than 150 µL of binding buffer. The tubes were gently tapped to mix every 15 min for 1 h total. After the incubation the beads were spun down and the supernatant discarded. The beads were washed 3 x 200 µL. The 4th wash of 100 µL was saved to check for residual protein. Loading dye was added directly to beads and boiled for 5 min. Following Run SDS PAGE and protein transfer the blot was stained with ponceau S solution. The blot was then washed clean with TBST and blotted for target proteins.

### Sample preparation for ubiquitylome and global proteome profiling

Soluble protein lysate (10 mg per condition) in denaturing buffer (8 M Urea, 20 mM HEPES, pH 8.0) was reduced (5 mM dithiothreitol (DTT), 45 min at 37 °C), alkylated (15 mM iodoacetamide (IAA), 20 min at room temperature in the dark), and quenched (5 mM DTT, 15 min at room temperature in the dark). Proteins were pelleted by chloroform-methanol precipitation. The resulting pellet was re-suspended in a denaturing buffer, diluted to 4M Urea, and digested for 4 h at 37 °C with lysyl-endopeptidase (Wako) at an enzyme-to-protein ratio of 1:100. The sample was further diluted to 1.3 M Urea and subjected to overnight enzymatic digestion at 37 °C with sequencing grade trypsin (Promega, enzyme : protein ratio = 1:50). Resultant peptides were acidified with 20% Trifluoroacetic acid (TFA, 1% final concentration), and an insoluble fraction was removed by centrifugation at 18,000 x g for 15 min. Peptide mixtures were desalted using a Sep-Pak C18 column (Waters). Eluted peptides from each treatment group (∼200 µg) were lyophilized and reserved for global proteome abundance analysis. The remaining eluted peptides were lyophilized and used for di-glycine (KGG) analysis.

For global proteome profiling, 100 µg of peptides from each sample were dissolved in 100 mM HEPES pH 8.0 (1 mg/mL). Isobaric labeling was performed using TMTPro16-plex reagents (Thermo Fisher). Each unit (0.5 mg) of TMT reagent was allowed to reach room temperature immediately before use, spun down on a benchtop centrifuge, and re-suspended with occasional vortexing in 20 µL anhydrous acetonitrile (ACN) prior to mixing with peptides (18% final ACN concentration). After incubation at room temperature for 1 h, the reaction was quenched for 15 min with 20 µL of 5% hydroxylamine. Labeled peptides were combined in equimolar ratios and dried. The TMTpro-labeled sample was re-dissolved in 80 µL 0.1% TFA, centrifuged at 16,000 x g, and the supernatant was processed further. Offline high pH reversed-phase fractionation was performed on a 1100 HPLC system (Agilent) using an ammonium formate based buffer system. Peptides (400 µg) were loaded onto a 2.1 x 150 mm 3.5 µm 300 Extend-C18 Zorbax column (Agilent) and separated over a 75-min gradient from 5% to 85% ACN into 96 fractions (flowrate = 200 µL/min). The fractions were concatenated into 24 fractions, mixing different parts of the gradient to produce samples that would be orthogonal to downstream low pH reversed phase LC-MS/MS. Fractions were dried and desalted using C18 stage-tips as previously described (Rappsilber et al., 2007). Peptides were lyophilized and re-suspended in 10 µL Buffer A (2% ACN, 0.1% formic acid) for LC-MS/MS analysis.

For ubiquitylome quantitation of KGG peptides, lyophilized peptides were reconstituted in 1X detergent containing IAP buffer (Cell Signaling Technology) for immunoaffinity enrichment. KGG peptide enrichment was performed at 4 °C on a MEA2 automated purification system (Phynexus) using 1 mL Phytips (Phynexus) packed with 20 µL ProPlus resin coupled to 200 µg of anti-KGG (Cell Signaling Technology) antibody. Phytip columns were equilibrated for 2 cycles (1 cycle = aspiration and dispensing, 0.9 mL, 0.5 mL/min) with 1 mL 1X IAP buffer prior to contact with peptides. Phytip columns were incubated with peptides for 16 cycles of capture, followed by 6 cycles of wash, 2X with 1 mL 1X IAP buffer and 4X with 1 mL water. Captured peptides were eluted with 60 µL 0.15% TFA in 8 cycles where the volume aspirated/dispensed was adjusted to 60 µL. Enriched ubiquitinated peptides were prepared as previously described (Rose et al., 2016). Labeled peptides were combined, dried, and re-solubilized in 0.15% TFA for high pH reversed-phase fractionation using a commercially available kit (Thermo Fisher). Fractionation was performed according to the manufacturer’s protocol with a modified elution scheme where 11 fractions were collected (F1: 13.5% ACN, F2: 15% ACN, F3: 16.25% ACN, F4: 17.5 ACN,F5: 20% ACN, F6: 21.5% ACN, F7: 22.5% ACN, F8: 23.75% ACN, F9: 25% ACN, F10: 27.5% ACN and F11: 30% ACN) and then combined into 6 fractions (F1+F6, F2+F7, F8, F3+F9, F4+F10, F5+F11). Peptides were lyophilized and re-suspended in 10 µL Buffer A for LC-MS/MS analysis.

### Mass spectrometry analysis for ubiquitylome and global proteome profiling

For global proteome and ubiquitylome quantitation of KGG peptides, LC-MS/MS analysis was performed on an Orbitrap Eclipse mass spectrometer (Thermo Fisher) coupled to a Dionex Ultimate 3000 RSLC (Thermo Fisher) employing a 25 cm IonOpticks Aurora Series column (IonOpticks, Parkville, Australia) with a gradient of 2% to 30% buffer B (98% ACN, 2% H2O with 0.1% FA, flow rate = 300 nL/min). Global proteome samples were analyzed with a total run time of 95 min and ubiquitylome samples were analyzed with a total run time of 180 min. For all samples, the Orbitrap Eclipse with FAIMS Pro DUO of -40, -60CV (proteome) or -40, -60, and -75CV (KGG) collected FTMS1 scans at 120,000 resolution with an AGC target of 1 x 106 and a maximum injection time of 50 ms. FTMS2 scans on precursors with charge states of 3-6 were collected at 15,000 resolution with CID fragmentation at a normalized collision energy of 30%, an AGC target of 2 x 104 (proteome) or 2 x 105 (KGG), a max injection time of 100 ms (proteome) or 200 ms (KGG). Real-time database search (RTS) was performed prior to acquisition of MS3 spectra using InSeqAPI software. The following RTS parameters were used for global proteome analysis: Uniprot human database January 2023 version, including 229,311 Swissprot sequences of canonical and protein isoforms, plus common contaminants and decoys; static modifications included Cys carbamidomethylation (+57.0215), Lys and n-term TMTPro (+304.207146); variable modifications included Met oxidation (+15.9949) and Tyr TMTPro (+304.207146). The following RTS parameters were used for KGG analysis: Uniprot human database August 2021 version, including 218,136 Swissprot sequences of canonical and protein isoforms, plus common contaminants and decoys; static modifications included Cys carbamidomethylation (+57.0215), Lys and n-term TMTPro (+304.207146); variable modifications included Met oxidation (+15.9949), Lys di-glycine (+114.042927) and Tyr TMTPro (+304.207146). All raw data has been deposited in the MassIVE repository under accession number: MSV000096813

Offline search was performed using comet v.2019.01 with parameters matched to the RTS search. Peptide FDR was filtered to <2% using Linear Discriminator Algorithm. TMT reporter ions produced by the TMT tags were quantified with Mojave in-house software package by calculating the highest peak within 20 ppm of theoretical reporter mass windows and correcting for isotope purities.

Quantification and statistical testing of global and KGG proteomics data were performed by MSstatsPTM v1.2.4 R package^45^. Peptide spectral matches (PSMs) were filtered out if they were (1) from decoy proteins; (2) from peptides with length less than 7; (3) with isolation specificity less than 50%; (4) with summed reporter ion intensity (across all channels) lower than 30,000. Multiple fractions from the same TMT mixture were combined in MSstatsTMT v2.0.1^46^. In particular, if the same peptide ion was identified in multiple fractions, only the single fraction with the highest maximal reporter ion intensity was kept. Global median normalization was carried out to reduce the systematic bias between channels. The normalized reporter ion intensities of all the peptide ions mapped to a protein were summarized into a single protein level intensity in each channel and TMT mixture by MSstats v4.0.1^47^. For quantification of relative lysine ubiquitination (KGG) levels, reporter ion intensities of all peptide ions mapped to a K-GG site were summarized into a single site level intensity in each TMT channel using MSstats. The differential abundance analysis between conditions (log2(fold change)) was performed in MSstatsTMT based on a linear mixed effects model per protein or KGG site. The inference procedure was adjusted by applying an empirical Bayes shrinkage. To test the two-sided null hypothesis of no changes in abundance, the model-based test statistics were compared to the Student t-test distribution with the degrees of freedom appropriate for each protein or site. The resulting p values were adjusted to control the FDR with the method by Benjamini-Hochberg. As a final step, changes in KGG levels were adjusted to the changes in total protein levels by MSstatsPTM.

### CRISPR KO resistance screening

A synthetic sgRNA library targeting 1384 genes related to ubiquitination was screened to identify genes required for compound **4** to degrade SMARCA2 protein. NCI-H1944 cells stably expressing S. pyogenes Cas9 from vector pLenti6.3 were arrayed into 384-well plates (PhenoPlate, PerkinElmer) at 850 cells per well and reverse transfected with pools of 3 sgRNAs targeting each gene (Synthego Corporation) at 10 nM concentration using Lipofectamine RNAiMAX reagent (0.05 µl per well in 41 µl total volume; Invitrogen). Testing cullin genes in advance of the screen identified CUL1 as a positive control for which sgRNAs blocked compound **4** -mediated SMARCA2 degradation. Each screening plate contained quality control wells for increased or decreased SMARCA2 staining (CUL1 or SMARCA2 sgRNAs, respectively), decreased cell proliferation (PLK1 sgRNAs), Cas9-mediated DNA damage (olfactory receptor OR5M9 and OR6N1sgRNAs), and no sgRNA (with and without compound **4**), and was screened with 3 or 4 replicates. After 4 days of transfection, either 0.5 µM of compound **4** or control 0.005% DMSO was added for 18 h, and cells were fixed with 4% paraformaldehyde (ThermoFisher Scientific) in PBS for 15 min at room temperature. Fixed cells were washed with PBS containing 0.5 mM MgCl2 and 0.5 mM CaCl2 (PBSCM) three times for 3 min each, permeabilized with 0.1% Triton X-100 in PBSCM (PBSCM-T) for 30 min, and then blocked with 3% BSA in PBSCM for 1 h. Cells were stained with anti-SMARCA2 antibody (D9E8B XP Rabbit mAb #11966, Cell Signaling Technology) diluted 1:2000 in blocking buffer at 4 °C for 16 h, and then washed with PBSCM-T three times for 3 min each. Cells were incubated in anti-rabbit IgG secondary antibody (Alexa Fluor 647 Conjugate #4414, Cell Signaling Technology) diluted 1:5000 and 2 µM Hoechst 33342 (ThermoFisher Scientific) for 1 h at room temperature, washed with PBSCM-T three times for 3 min each, and then resuspended in PBS. Confocal fluorescence microscope images were acquired with the ImageXpress Micro Confocal High-Content Imaging System (Molecular Devices) using a 10x objective and processed with the MetaXpress High-Content Image Acquisition and Analysis Software for SMARCA2 immunofluorescence integrated intensity and nuclear counts.

A robust z-score transformation was performed for screening hit identification within each plate to standardize SMARCA2 immunofluorescence intensity values across plates; the location and scale factors were estimated by taking the median and median absolute deviation (MAD) of the intensity values across all experimental genes. A differential analysis was then conducted between conditions using the limma R package on the z-score transformed values^48^. p-values were corrected for multiple comparisons using the Benjamin-Hochberg procedure, resulting in FDR-corrected p-values. 25 novel screen hit genes for which sgRNAs block SMARCA2 degradation by monovalent degrader **4** were selected by the criteria of a difference in z-scores greater than 3 and -log10(FDR-adjusted p) greater than 1.3. Five of these screen hits, including FBXO22, were further distinguished as compound **4** -selective in their dependency for SMARCA2 degradation based on their median SMARCA2 immunofluorescence intensity values not being above that of negative control OR5M9 and OR6N1 sgRNAs for DMSO-treated cells.

### SMARCA2 NanoBRET^®^ Assay

The SMARCA2 NanoBRET^®^ Target Engagement Assay assessed the apparent binding affinity of test compounds for SMARCA2 in permeabilized HEK293 cells by competitive displacement of a tracer. The tracer, which was created with a SMARCA2 ligand and NanoBRET^®^ 590 SE Dye, reversibly bound to a SMARCA2-NanoLuc^®^ (C-terminal) fusion protein transiently expressed in HEK293 cells. Test compounds and DMSO were transferred to the assay plate (384-Well White, Non-Binding Corning Assay Plate (Corning-3574) using an Echo 555 Liquid Handler (Labcyte) in order to prepare a titration series. Maximum Signal control wells consisted of 80 nL DMSO-only treated wells. Minimum Signal control wells contained 10 µM parental unlabeled SMARCA2 ligand. Background Signal control wells were prepared without tracer for background correction steps. DMSO was backfilled to a final volume of 80 nL as required (0.2% DMSO in a 40 µL reaction). 200 nL per well of tracer in DMSO were transferred into each well using the Echo 555 for a final concentration of 0.25 µM tracer. HEK293 cells were cultured in DMEM High Glucose with Pyruvate, 10% fetal bovine serum, 1% Gibco™ Antibiotic-Antimycotic (Thermo Fisher 15240062). Bulk transfection was performed with FuGENE® HD (40 µL : 500 µL of Transfection Reagent:DNA mixture in Opti-MEM (Thermo Fisher 11058-021) overnight. Digitonin (Sigma D5628-1G) was added to cells right before plating for a concentration of 10 µg/mL. 10,000 cells per well in a 40 µL well volume were dispensed using the Multidrop™ Combi Reagent Dispenser and incubated for 15 min. 3x Complete Substrate plus Inhibitor Solution was prepared in Opti-MEM (consisted of a 1:166 dilution of NanoBRET® Nano-Glo® Substrate), and 20 µL was dispensed into each well of the 384-well plate. Plates were read using a PerkinElmer Envision Reader (model 2104-0020) after 2-minute incubation at room temperature. The raw BRET ratio values were calculated by dividing the acceptor emission value (610 nm) by the donor emission value (460 nm) for each sample. Raw data was normalized and scaled to the DMSO control column (Maximum Signal) on a per row basis to account for decrease of substrate during plate read time and was then background corrected with the no tracer condition (Background Signal) before fitting normalized BRET (%) compound dose-responses with software such as GraphPad Prism. The IC_50_ was then converted to an apparent compound KD using the Cheng-Prusoff correction by dividing the IC_50_ by a constant. The constant was separately determined by tracer titration with digitonin (data not shown).

### SMARCA2/FBXO22 NanoBRET® Assay

FBXO22 knockout HCC515 cells were plated in 6-well plates (800,000 cells/well) overnight and subsequently transiently transfected with SMARCA2-NanoLuc^®^ and FBXO22-HaloTag^®^ (Promega, FHC25469) at 1:20 ratio of NanoLuc® to Halotag^®^ protein using FuGENE HD (Promega, E2311). On the following day, cells were harvested and replated at 25,000 cells/well in a 96-well assay plate in the presence or absence of HaloTag^®^ NanoBRET**^®^** 618 Ligand (Promega, G9801). Cells were incubated overnight and on the next day treated with DMSO or varying concentrations of **G-6599** and control PROTAC, A947. Cells were pretreated for 1 h with 10 µM MLN4924 and maintained in MLN4924 for the duration of the experiment. After 24 h treatment, the NanoBRET**®** Nano-Glo**^®^** Substate (Promega, N1571) was added in Opti-MEM (Thermo Fisher Scientific, 31985062). The donor (460 nM) and acceptor (618 nM) emissions were measured on the NanoBRET**®** Nano-Glo**®** Detection System (Promega, N1661) 10 min after adding the substrate. The raw NanoBRET**^®^**ratio was calculated by dividing the acceptor emission value by the donor emission value. That value was then multiplied by 1,000 to convert to milliBRET units (mBU). The corrected NanoBRET**^®^** ratio was calculated by subtracting the raw NanoBRET**^®^** ratio (mBU) of the no NanoBRET**^®^** 618 Ligand control from the raw NanoBRET**^®^**ratio (mBU) of the same treatment with NanoBRET**^®^** 618 ligand.

#### HDX-MS

Experiments were conducted using an extended, parallel-arm LEAP HDX automation system (Trajan Scientific, Chronos software) interfaced with a Thermo Ultimate 3000 HPLC and Exploris 480 mass spectrometer. To initiate deuterium labeling, a 5 µL aliquot of 10 µM of FBXO22 material, with or without 100 µM SMARCA2 (BRM) or 100 µM **G-6599** was combined with 60 µL of D_2_O-based, 20 mM HEPES-Cl buffer, pDread 7.1, and 150 mM NaCl. After fixed periods of labeling time at 5 °C (60, 294, 1440 min for SMARCA2 experiments, or 1.0, 1.98, 10.31, 53.52, 277.58, and 1440.0 (apo FBXO22) or 2280.0 (bound) min for SMARCA2+**G-6599** experiments), 55 µL of this mixture was combined with 55 µL of an H_2_O buffer containing 4 M GdmCl, 1 M glycine, pH 2.50, 95 µL of this mixture then injected into a 0 °C (+/- 1°) temperature controlled chromatography chamber the sample was digested, online, with a 50:50 Fungal protease / Pepsin column (NovaBioAssays) and then loaded onto a C8 trap column (Waters, Acquity UPLC BEH C8 VanGuard Pre-column, 130 Å, 1.7 µm, 2.1 mm X 5 mm) and then washed for approximately 3 min (150 µL/min, mobile phase A) before being separated by acetonitrile gradient using a C18 analytical column (Waters, ACQUITY UPLC BEH C18 Column,130 Å, 1.7 µm, 1 mm X 50 mm). Peptides are introduced into the gas phase by electro-spray ionization (Spray voltage Static, Positive mode - 3400V, Sheath Gas 15, Aux Gas 10, Sweep Gas 1, Ion Transfer Tube Temperature 220 ℃, Vaporizer Temperature 75 ℃). Unless otherwise specified, all data was collected in MS-only mode (120K Hz). Each timepoint was sampled in triplicate.

The same procedure was used, without deuterium, and analyzed by tandem-MS (60k Hz resolution (MS1) 30k Hz resolution (MS2)) to identify peptides. Further, a non-deuterated control was measured on the day of the labeling experiments, using the instrument parameters described above, for retention time referencing. Protein Metrics Byonic and Byologic software was used for peptide identification (FDR2D filter of ≤ 0.0001) followed by ExMS 2.0 to provide XIC’s and deuterium uptake levels for each peptide across the sampled timepoints using recommended settings^49^. The empirical method was employed to extract protection factors from each peptide^50^.

For experiments involving FBXO22+SMARCA2, 234 peptides cover 92.6% of the FBXO22 sequence, with a mean redundancy of 7.7 unique peptides per residue, and mean peptide length of 13.1 amino acids. For experiments involving FBXO22+SMARCA2+**G-6599**, 241 peptides cover 93.5% of the FBXO22 sequence, with a mean redundancy of 6.6 unique peptides per residue, and mean peptide length of 12.4 amino acids (optional coverage map figures). Per reporting standards suggested previously^51^: all peptides are available for inspection (See Supplementary Figures 1, 2) and the PFs extracted and used in the analysis along with information about each peptide from which the PF was derived are reported in Supplementary Table 6. All processed peptides were manually confirmed to be consistent with other overlapping peptides. All raw data has been deposited in the MassIVE repository under accession number: MSV000097140

Software used for HX experiments:

EXMS 2.0 Version 03NOV2018

Protein Metrics Byonic Version 4.5.2

Protein Metrics Byologic Version 4.5-53 x64

Chronos Version 5. 1. 10

MSConvertGUI Version 3.0.22027-235693e

### Covalent labeling mass spectrometry

Mixtures containing 1 µM FBXO22 WT and 20 µM compound (**G-6599**) (+/- 1 µM SMARCA2) with 2.5% (v/v) DMSO were incubated at room temperature for 4 h in PBS buffer [137 mM NaCl, 10 mM phosphate, and 2.7 mM KCl (pH 7.4)]. The reaction was quenched by adding formic acid to 1%. An aliquot (5 µL) of the sample was submitted to LC-MS. Accurate mass data were obtained on an Agilent 6230 time-of-flight mass spectrometer in positive ion mode equipped with a dual Agilent Jet Spray (AJS) ion source in positive ionization mode, an autosampler (held at 6 °C), and an Agilent 1290 binary pump. The sample was loaded onto a Waters BEH C4 column (2.1 mm × 150 mm, 300 Å, 1.7 µm) held at 45 °C. Mobile phase A was a solution of 0.1% formic acid in water. Mobile phase B was a solution of 0.1% formic acid in acetonitrile. The flow rate was set to 500 µL/min. The gradient used was 5% B for 0.8 min, ramping linearly to 50% B from 0.8 to 5 min, holding at 80% B from 5.2 to 5.5 min, and returning to 5% B at 5.6 min, and the column was allowed to equilibrate for 0.4 min before the next injection was initiated. The spectra were acquired from 550 to 3200 Da using a gas temperature of 350 °C, a gas flow of 10 L/min, and the nebulizer gas at 35 psi. The following voltages were used: capillary, 5000 V; fragmentor, 250 V; skimmer, 100 V; and octapole RF peak, 750 V. Spectra were acquired at a rate of one spectrum per second. The data were processed using MassHunter software.

### Peptide mapping mass spectrometry

Recombinant FBXO22/SKP1 and SMARCA2 proteins were diluted to a final concentration of 2 µM in phosphate-buffered saline (PBS) and mixed in equal volumes to obtain a protein mixture containing 2 µM FBXO22/SKP1 and 2 µM SMARCA2. Compound **G-6599** was diluted in PBS from a 10 mM stock in DMSO to a final concentration of 100 µM. An equal volume of the protein mixture and compound solution were combined, resulting in final concentrations of 2 µM proteins and 50 µM compound in a total volume of 60 µL. The reaction was incubated at room temperature for 4 h.

Reduction was performed by adding dithiothreitol (DTT) and cysteine to final concentrations of 5 mM and 1 mM, respectively, followed by incubation at 37 °C for 30 min. Urea was then added to a final concentration of 6 M by adding 75 µL of 10 M urea solution. Alkylation was carried out by adding iodoacetamide to a final concentration of 20 mM and incubating in the dark at room temperature for 30 min. The solution was diluted with 250 µL PBS to reduce the urea concentration to 2 M.

The sample was divided into two equal aliquots, each containing approximately 2 µg of protein in 180 µL. Protease digestion was performed using either trypsin/LysC or chymotrypsin (Promega) at a protease-to-protein ratio of 1:25 (w/w). Proteases were diluted to 100 ng/µL, and 0.8 µL (80 ng) was added to each aliquot. Digestion proceeded at 37 °C for 16 h and digestion was quenched by adding formic acid to a final concentration of 1% and incubating at room temperature for 10 min.

Peptides were desalted using C18 tips (Thermo Fisher), concentrated by speed vacuum centrifugation, and resuspended in 18 µL of 0.1% formic acid. For LC-MS/MS analysis, 6 µL (approximately 660 ng) of the peptide solution was loaded onto the chromatographic system. Samples were pre-concentrated on a PepMap C18 column (Thermo Scientific, 300 µm × 5 mm, 5 µm, 100 Å) and desalted for 4 min. Chromatographic separation was performed using an EASY-Spray PepMap C18 column (75 µm × 150 mm, 2 µm, 100 Å) on a Vanquish Horizon UHPLC system (Thermo Scientific). The mobile phase consisted of solvent A (water with 0.1% [v/v] formic acid) and solvent B (100% [v/v] acetonitrile with 0.1% [v/v] formic acid). Peptides were eluted at a nominal flow rate of 400 µL/min, with a T-shaped microfluidic junction incorporated in the flow path prior to the pre-concentration and analytical columns. This junction further split and reduced the flow rate during peptide elution, resulting in an estimated post-junction flow rate of approximately 1 µL/min in peptide elution. The gradient was 1% B to 40% in 20 min, increased to 80% in 0.1 min, held at 80 % for 4.9 min, return to 1% B in 0.1 min and equilibrated at 1% B for 7 min. The eluent was directed to an EASY-Spray source coupled to an Orbitrap Exploris 480 mass spectrometer (Thermo Fisher) for detection. The spray voltage was set to 1900 V, with a capillary temperature of 275°C. In the data-dependent mode, the 6 most-abundant ions were subjected to higher energy collisional dissociation (HCD) with an isolation window of m/z 2 and a normalized collision energy of 26%.

### Metabolite Identification in HCC515 Cell and Media Incubations

The test compound (Compound **5** or **G-6599**) was incubated at 1 µM in 50 µL of HC5515 cells or basal media with or without 10% FBS for 0, 1, and 4 h. The cell density was approximately 0.5 x 10^6^ cells/mL. Incubations were quenched with 3 volumes of ACN + 0.1% formic acid with 50 nM propranolol as an internal standard. The samples were mixed, centrifuged, and the supernatant was concentrated under N_2_ gas. The resulting samples were analyzed by LC-MS using a Vanquish HPLC system and Orbitrap Fusion Lumos mass spectrometer (Thermo Fisher Scientific). The chromatography used a Luna Omega Polar C18 column (1.6 µm, 2.1 × 100 mm; Phenomenex) at 40 °C with a flow rate of 0.4 mL/min. Mobile phase A was 0.1% formic acid in water, and mobile phase B was 0.1% formic acid in acetonitrile. The column was initially held at 2% B for 1 min, increased to 30% B over 11 min, and then increased to 95% B over 0.5 min, where it was maintained for 1 min before returning to 5% B over 0.5 min for re-equilibration. Mass spectrometry data was acquired using positive ion mode and a full scan-data-dependent MS2 mode.

Metabolite identification was performed by processing and analyzing the LC-MS data using MassMetaSite and ONIRO (Molecular Discovery). Metabolite structures were proposed based on analysis of the high-resolution accurate mass data and the MS2 fragmentation data, and all analytes were determined to have accurate masses within 5 ppm. For each sample, the relative mass spectrometry abundances of the parent compound and its metabolites and impurities were calculated as the extracted ion chromatogram (XIC) peak area, including any relevant charge states, relative to the sum of the XIC peak areas for all reported analytes in that sample.

### Cellular thermal stability shift assay (CETSA)

HCC515 cells were plated in 10 cm dishes and grown to ∼95% confluency. Cells were subsequently washed 2x w/PBS and harvested by scraping in KB buffer (25 mM Tris HCl, pH 7.5, 2 mM DTT, 10 mM MgCl_2_, w/ protease and phosphatase inhibitors). Samples were lysed by snap freezing 3x in liquid N_2_ and subsequently incubated on ice for 10 min. Lysates were cleared by centrifugation at 20,000 g for 20 min at 4 °C and supernatants were transferred to clean tubes. Lysates were sonicated briefly and concentrations were adjusted to 1.5 mg/mL with KB buffer. Following addition of 10 µM neddylation inhibitor for 10 min, compounds or DMSO control was added and lysates were incubated at room temperature for 30 min. Lysates were aliquoted to PCR strip tubes (50 µl) and heated at 38 °C to 71 °C (3 degree increments) for 3 min in a PCR machine (Applied Biosystems Veriti Thermal Cycler). Following cooling, tubes were centrifuged at 20,000 g for 20 min at 4 °C and transferred to clean tubes. Sample buffer was added prior to boiling lysates and running on SDS-PAGE gels.

## Acknowledgements

The authors would like to thank Robert Blake for providing the SMARCA2-NanoLuc® construct, Pirunthan Perampalam for his contribution to the analysis of the CRISPR screen, Megan Flynn for providing SMARCA2 bromodomain protein and Szymon Juszkiewicz and Peter Ung for critical discussions related to this work. Liting Dong, Yexia Zhang, Yongsheng Cheng and Zhijuan Cai from WuXi AppTec chemistry and parallel synthesis teams for the synthesis of SMARCA2/A4 monovalent degraders. Kewei Xu, Yuhui Zhou and Nazareo Gonzalez from Genentech Analytical Research for assistance with compound QC.

---

**Extended Data Fig 1.**
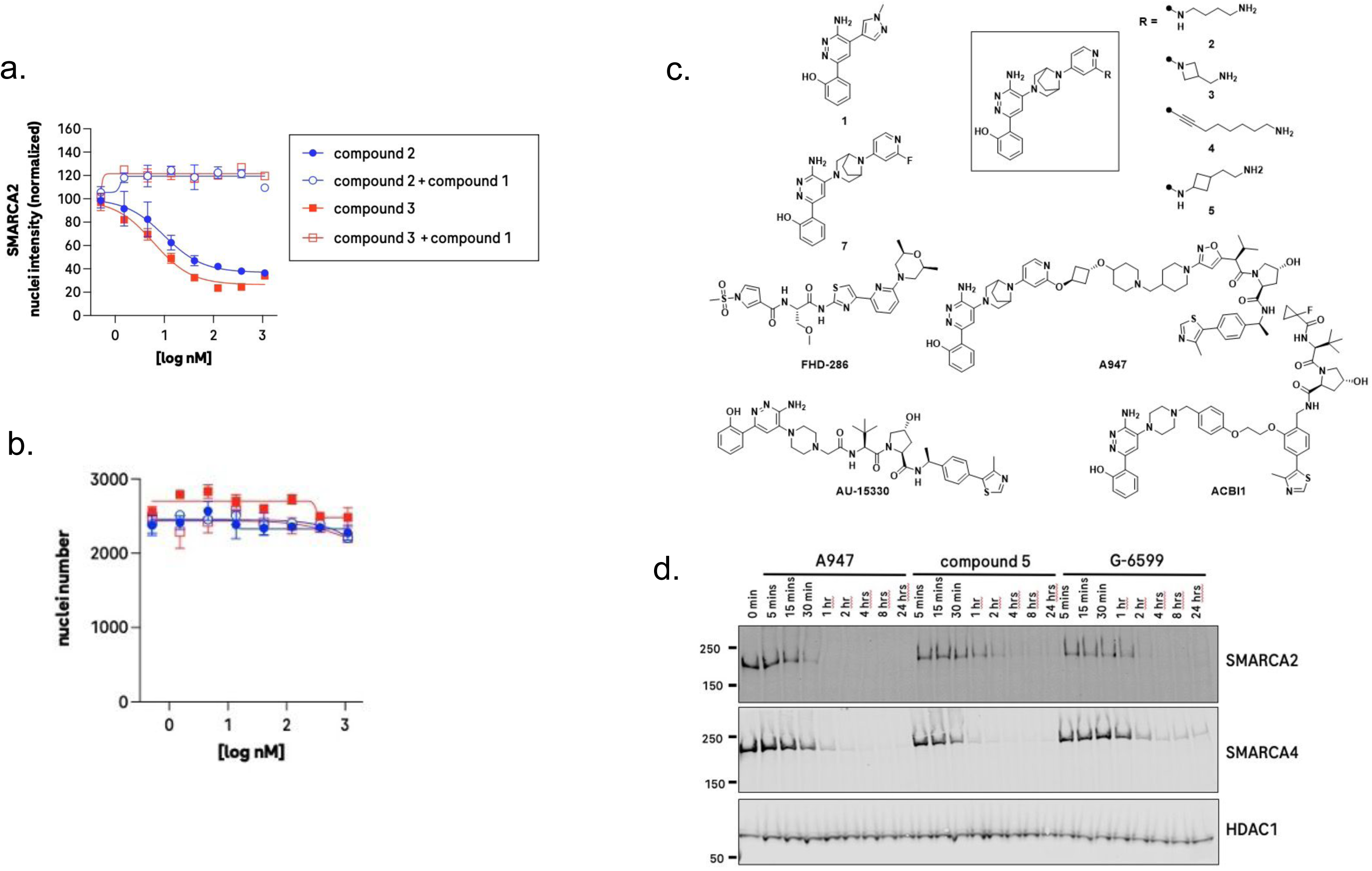
**a,** Dose-response curves of compound **2** and **3** on SMARCA2 levels detected by immunofluorescence in SW1573 cells in the presence/absence of 50 uM parent SMARCA2/A4 ligand, compound **1**. **b,** Nuclei count following treatment of SW1573 cells with dose-response of compounds **2** and **3**. **c,** Chemical structures of additional molecules utilized in this manuscript. **d,** Immunoblot analysis of the SMARCA2 and SMARCA4 in SW1573 cells following a timecourse of treatment with monovalent degraders or PROTAC control (A947). HDAC1 serves as a loading control.

**Extended Data Fig. 2.**
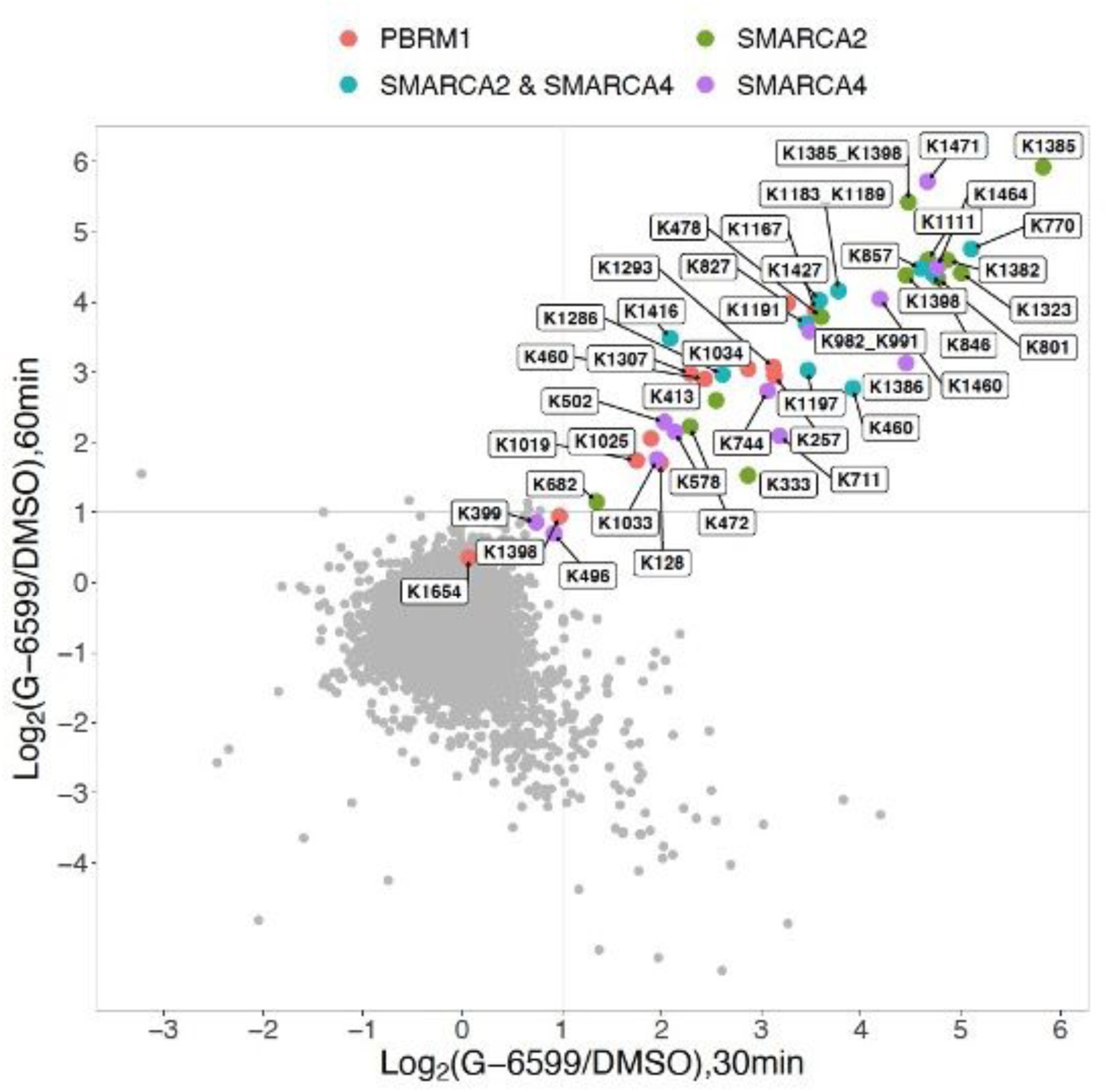
Global ubiquitylome changes assessed by di-glycine remnant mass spectrometry profiling in SW1573 cells following 30 and 60 min treatment with 100 nM **G-6599**. Peptides mapping to SMARCA2 (UniProtKB P-51531-2), SMARCA4 (UniProtKB P-51532-1), PBRM1 (UniprotKB Q86U86) or SMARCA2/A4 shared peptides are highlighted in color. n=9195 unique ubiquitinated sites were identified and presented as log2 fold change relative to DMSO control cultures.

**Extended Data Fig. 3.**
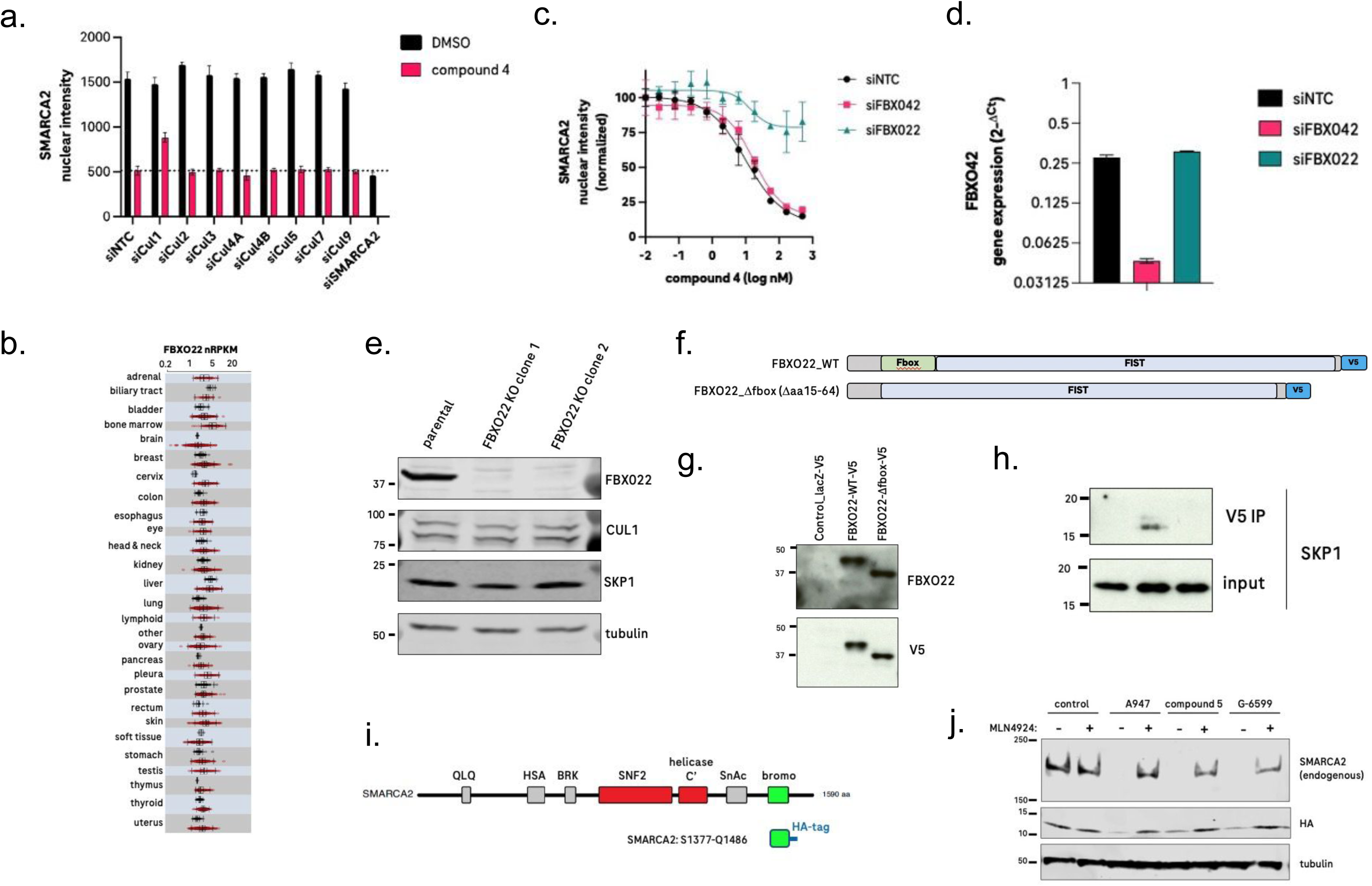
**a,** Immunofluorescent-based detection of SMARCA2 in HCC515 cells transfected with siRNAs targeting specific cullins or SMARCA2 following 8 h treatment with compound **4** (200 nM) or DMSO control. Data are presented as mean ± s.d from quadruplicate cultures. **b,** mRNA expression of FBXO22 measured using RNASeq in human tissues collected by TCGA. Gene expression is displayed in normalized Reads Per Kilobase of exon model per Million mapped reads (rpkm). Black and red dots refer to normal and tumor specimens, respectively. **c**, Effect of siRNA-mediated knockdown of FBXO22, or a control E3 ligase, FBXO42, on SMARCA2 levels in HCC515 cells following treatment with varying concentrations of compound **4**. A non-targeting control siRNA (siNTC) served as a control **d,** mRNA expression values of FBXO42 determined by Taqman following siRNA treatment of HCC515 cells. Data are presented as mean ± s.d from triplicate cultures. **e,** Immunoblots demonstrating FBXO22 loss in FBXO22 knockout clones relative to parental HCC515 cells. Tubulin serves as a loading control. No change in Cul1 or SKP1 expression in FBXO22 knockout cells was detected. **f,** Domain plots of V5-tagged FBXO22 constructs. **g,** Immunoblots of FBXO22 knockout HCC515 cells stably reconstituted with V5-tagged WT or ΔFbox FBXO22. Cells expressing lacZ-V5 served as a control. **h,** SKP1 immunoblots of input lysate or anti-V5 immunoprecipitates from cells in (g). **i,** Depiction of SMARCA2 structure and HA-tagged SMARCA2 bromodomain construct. **j,** Immunoblot analysis of HCC515 cells transfected with HA-tagged SMARCA2 bromodomain following overnight treatment with 200 nM of the respective compounds or DMSO control. When specified, cells were pretreated for 1 h with 6 µM MLN4924 and maintained in MLN4924 for the duration of the experiment.

**Extended Data Fig. 4.**
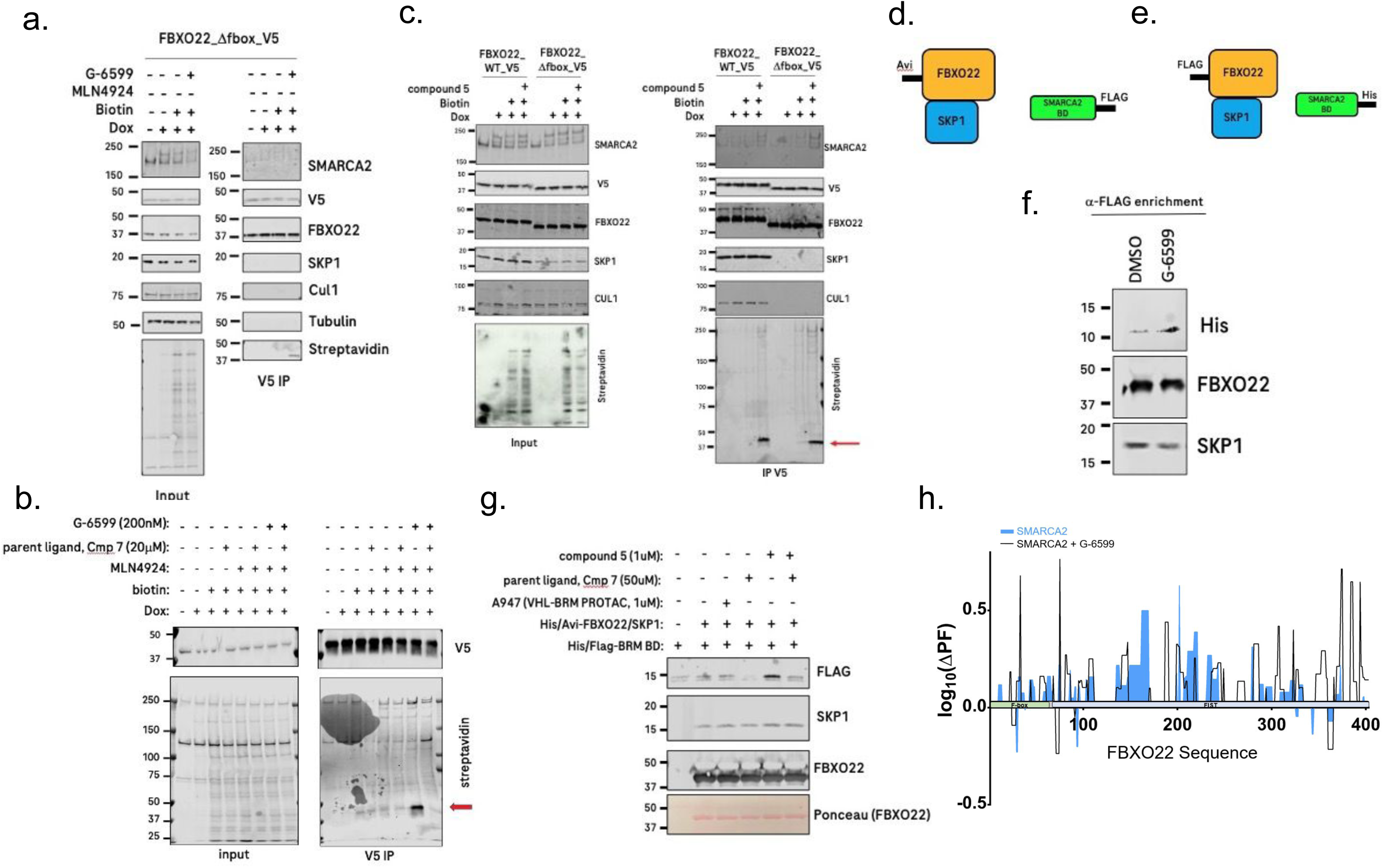
**a,** FBXO22 knockout HCC515 cells stably expressing V5-tagged ΔFbox-FBXO22 and doxycycline-inducible SMARCA2-TurboID were pre-treated for 72 h with 1 µg/mL doxycycline followed by a 1 h treatment with 6 µM MLN4924 and subsequent addition of 200 nM **G-6599** and 50 µM biotin overnight. Biotinylated proteins were detected by streptavidin Western blotting of input and anti-V5 immunoprecipitates. Tubulin served as a negative control protein in V5 immunoprecipitates. **b,** FBXO22 knockout HCC515 cells stably expressing V5-tagged FBXO22 and doxycycline-inducible SMARCA2-TurboID were treated as in (a), but included pre-treatment with 20 µM parent ligand to compete with **G-6599**. Biotinylated proteins were detected by streptavidin Western blotting of input and anti-V5 immunoprecipitates. V5 immunoblots served as a control for V5-FBXO22 in immunoprecipitates. **c,** FBXO22 knockout HCC515 cells stably expressing V5-tagged WT or ΔFbox-FBXO22 and doxycycline-inducible SMARCA2-TurboID were treated as in (a), but treated with 200 nM compound **5**. **d, e** Depiction of recombinant tagged proteins used in Fig. 4f and Extended Data Fig. 4f. **f**, Recombinant FLAG-tagged protein immobilized on anti-FLAG resin was mixed for 1 h with His-tagged SMARCA2 protein (aa. S1377-Q1486) in DMSO or 1 µM **G-6599**. Precipitates were washed and immunoblotted for the respective proteins. **g**, Compound **5**-mediated co-immunoprecipitation of recombinant FBXO22 protein with recombinant SMARCA2 bromodomain can be competed by excess parental ligand in solution. Recombinant Avi-tagged FBXO22/SKP1 protein immobilized on anti-Avi resin was mixed for 1 h with FLAG-tagged SMARCA2 protein (aa. S1377-Q1486) in the respective conditions. Precipitates were washed and immunoblotted for the respective proteins. **h,** HDX-MS plot representing the log_10_ change in protection factor from deuterium uptake for SMARCA2 (light blue, full) and SMARCA2+**G-6599** (black line) in the presence of FBXO22 compared to FBXO22 alone aligned on FBX022 protein sequence.

**Extended Data Fig. 5.**
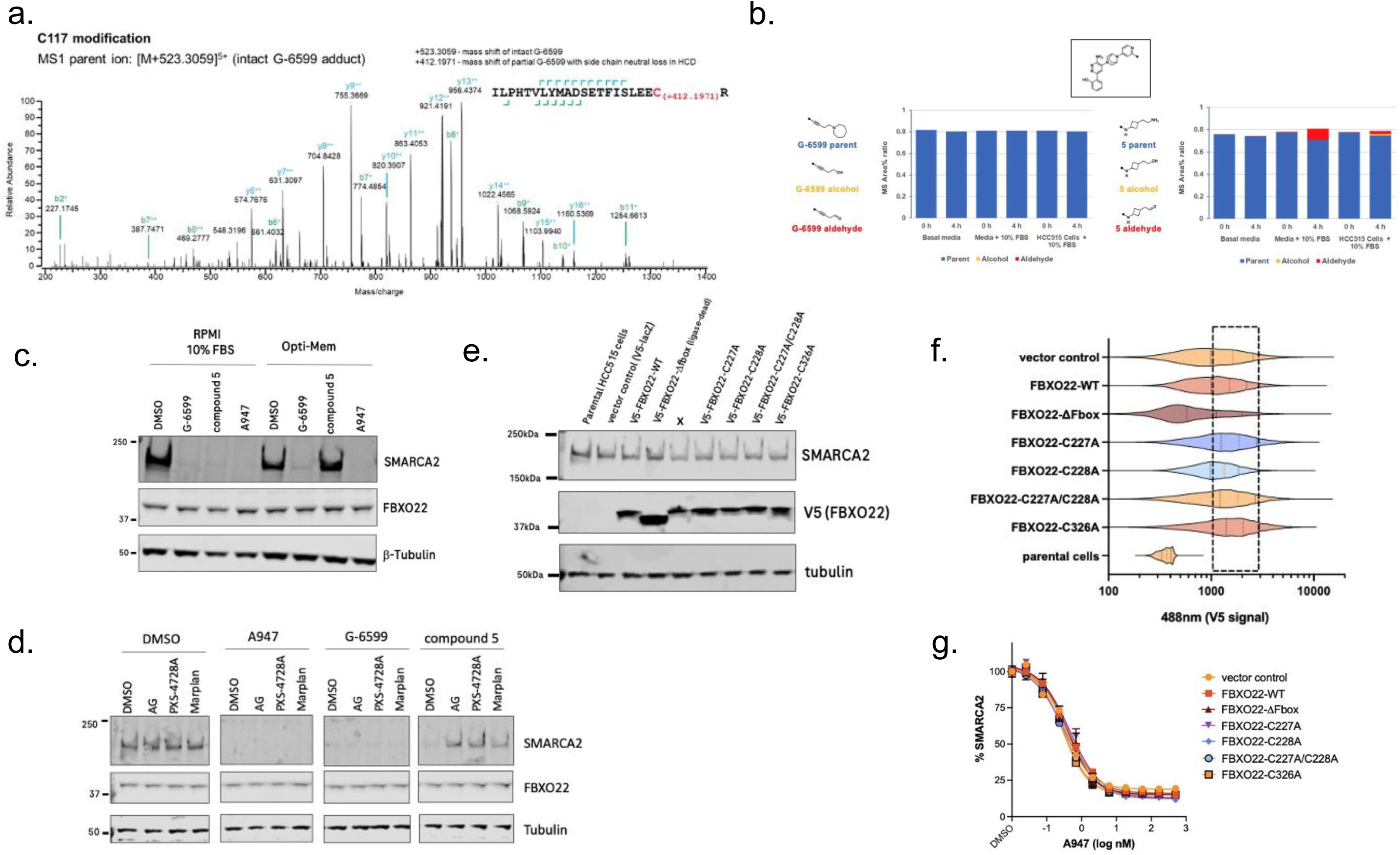
**a,** Identification of covalently modified FBXO22 peptides using mass spectrometry after 4 h treatment of 1 µM FBXO22/SKP1 and SMARCA2 bromodomain in the presence of 50 µM of **G-6599** for Cys117 modification. **b**, Metabolite identification using mass spectrometry after treatment of 1 µM of **G-6599** (left) or compound **5** (right) in cell media with or without 10% FBS or in the presence of HCC515 cells with 10% FBS for 0 h and 4 h. The mass area % ratios of the parent molecule (blue), the alcohol (yellow) and the aldehyde (red) metabolites are depicted. **c,** HCC515 cells were treated for 8 h with **G-6599** (200 nM), compound **5** (200 nM) or control PROTAC, A947 (100 nM) in RPMI media containing 10% FBS or serum-free Opti-MEM media prior to immunoblotting for the respective proteins. Tubulin serves as a loading control. **d,** HCC515 cells were pretreated for 30 min with the respective amine oxidase inhibitors (1 mM) prior to 4 h treatment with **G-6599** (200 nM), compound **5** (200 nM) or control PROTAC, A947 (100 nM) and were subsequently analyzed by immunblotting. Tubulin serves as a loading control. AG: aminoguanidine. **e,** Immunblot analysis of FBXO22-knockout HCC515 cells engineered to re-express wild-type or mutant, V5-tagged FBXO22. Tubulin serves as a loading control. **f,** Immunofluorescence detection of V5 protein levels across engineered cell lines in Ext. Data Fig 5e. For quantification of SMARCA2 levels upon treatment with **G-6599** in Fig. 5d, analysis was restricted to cells expressing comparable V5 levels, between 1000-3000 emission units (hashed line box). **g,** Effect of varying concentrations of control PROTAC, A947, on SMARCA2 levels in FBXO22 knockout HCC515 cells reconstituted with V5-tagged wild-type, or mutant FBXO22. Nuclear intensity of SMARCA2 within V5-positive cells is presented relative to DMSO control conditions

